# Narratives engage brain and body: bidirectional interactions during natural story listening

**DOI:** 10.1101/2023.01.31.526511

**Authors:** Jens Madsen, Lucas C. Parra

## Abstract

It is often said that the body and the mind are connected. Yet, direct evidence of a bidirectional link is elusive. We hypothesized a top-down effect of cognition on arousal, and predicted that auditory narratives will drive not only brain signals but also peripheral physiological signals. We find that auditory narratives entrained gaze variation, saccade initiation, pupil size, and heart rate. This is consistent with a top-down effect of cognition on autonomic function. We also hypothesized a bottom-up effect, whereby autonomic physiology affects arousal. Controlled breathing affected pupil size, and heart rate was entrained by controlled saccades. Additionally, fluctuations in heart rate preceded fluctuations of pupil size and brain signals. Gaze variation, pupil size and heart rate were all associated with anterior-central brain signals. Together this suggests bidirectional causal effects between peripheral autonomic function and central brain circuits involved in the control of arousal.

**Highlights:**

- Listening to narratives modulates eye movements.
- Heart rate fluctuations precede fluctuations in pupil size and anterior-central neural activity.
- Breathing modulates pupil size suggesting causal effect on central arousal.
- Rhythmic saccades can entrain heart beats.

**eTOC:** When we listen to a story our body is integrally involved in the experience. We provide evidence for a bidirectional and causal link between body and mind by analyzing brain signals, pupil size, heart rate and eye movements, while subjects listen to narratives and during interventions that control autonomic signals.

## Introduction

As we engage with the world we move our eyes scanning our surroundings. Fluctuations in lighting cause the pupil to contract or dilate^1^ and even in the absence of luminance changes our pupil fluctuates^2^. During task demand our autonomic system regulates dilation and constrictions of the pupil via sympathetic and parasympathetic activity^3^. A major hub of the autonomic system that regulates arousal is the locus coeruleus^4^. Its activity is tightly linked to pupil size^5^ and thus pupil size is often considered a marker of arousal^6^. These task-evoked pupil responses have been shown to entrain to task timing^7^, cognitive control^8^ and cognitive resource allocation in mental arithmetic tasks^9^. Interestingly, a number of cognitive factors modulate pupil size, such as affective processing^10–12^, cognitive effort^13^, cognitive control^14^, task engagement^15^, conscious processing^16^, attention^17, 18^ and more. This suggests a direct effect of cognition on arousal. Phasic fluctuations of LC neurons have been associated with task-related decision processes whereas tonic activity has been associated with disengagement from the current task and a search for alternative behaviors^19^.

The heart is also controlled by the autonomic nervous system, with sympathetic response increasing heart rate and parasympathetic activity decreasing it^20^. Physical activity has an obvious effect on heart rate^21^ as does arousal from sleep^22^. When we are resting, heart rate fluctuates with respiration, which is known as respiratory sinus arrhythmia^23^. Stressful mental effort can also increase heart rate^24^. Even subtle “arousal” due to emotional^25^ or surprising^26^ visual stimuli appears to accelerate individual heart beats. At a slower time scale of several heart beats, heart rate is positively correlated with cortical activity in fMRI predominantly in midline brain areas^27^. The overall strength of fluctuations -- measured as heart rate variability^28^ -- is affected by various cognitive factors^29–34^. We recently demonstrated that audiovisual narratives can reliably increase or decrease heart rate at different moments of the story, and thus synchronize heart rate across people^35^. This suggests that the cognitive processing of the narrative has an immediate effect on heart rate.

Eye movements have also been linked to arousal in a recent study^15^, which argues that a common subcortical signal controls pupil dilation as well as saccade initiation -- saccades are fast eye movements that redirect gaze to a new location of the visual field. We have recently found that engaging videos reliably modulate saccades rate (the number of saccades per second), but only if viewers are paying attention^36^. This suggests that cognitive processing of the visual stimulus modulates saccade rate. What is not yet known is whether a purely auditory narrative will also affect pupil size and eye movements, as we have observed for heart rate^35^. If heart rate fluctuated as a result of fluctuating arousal, we might expect that the pupil and saccades will be similarly driven by an auditory-only narrative. Here we hypothesized that the cognitive processing of an auditory narrative will drive not only heart rate, but also pupil size and eye movements.

To assess this we measured inter-subject correlation of these peripheral signals, as well as scalp potentials. Inter-subject correlation is an established metric to capture processing associated with dynamic natural stimuli such as films^37^ or auditory narratives^38, 39^. Scalp potentials are known to entrain to low-level features of continuous speech^40, 41^ as well as its semantics^42^ -- a phenomenon sometimes referred to as “speech tracking”^43^. To disambiguate semantic processing of the narrative from purely low-level auditory processing we will use attentive and distracted listening conditions^44^, and additionally analyze responses to semantic changes in the narrative, similar to previous work with scalp potentials^42^. With these experiments we aim to test whether there is a top-down effect of cognitive processing on pupil size and eye movements, without confounds of a visual stimulus or motor behavior, while controlling for the known effects of eye movements on pupil size^45, 46^.

To determine if there is a reverse, bottom-up effect we will use interventions that control these peripheral signals individually. We exploit the fact that heart rate is modulated by breathing^47^, increasing with inhalation and decreasing with exhalation. Pupil size is similarly affected by luminance^1^. Finally, we elicit controlled saccades by asking participants to follow a dot jumping on the screen. We hypothesize that these interventions will result in a modulation of the other peripheral signals. For instance, fluctuating heart rate (as a result of rhythmic breathing) might lead to rhythmic fluctuations of pupil size. Similarly, rhythmic saccades might affect heart rate. Finally, fluctuating lighting might affect heart rate. To our knowledge, none of these effects have previously been observed. Yet pupil size, heart rate, and eye movements are controlled by interconnected brainstem nuclei^15, 18, 29^. If we do find such interactions, it suggests that there are common brain nuclei centrally driving these signals. Importantly, a modulation of pupil size would suggest a modulation of cortical excitability, given the ample evidence that pupil size reflects bottom-up cortical activation during quiet wakefulness^48^, via the ascending arousal system^49, 50^. To summarize, we will explore a top-down effect of cognition on the peripheral signals of pupil size, eye movements and heart rate. And determine if bottom-up control of individual peripheral signals carries over to other peripheral signals in the absence of a cognitive drive.

## Results

We experimentally controlled four factors at play (Fig. 1A): Cognition through auditory narratives in an attentive (blue) and distracted condition (green), heart rate through controlled breathing (red), gaze position through controlled saccades or fixation cross (purple) and pupil size through controlled luminance (yellow). At the start we also included a rest condition (pink) to capture spontaneous brain-body interactions. In the distracted listening condition, participants were asked to perform a mental arithmetic task, which is meant to distract them from the narrative. To probe memory, towards the end of the experiment we asked participants questions to test recall of the content of the auditory narrative.

From previous work we know that listening to these autobiographical narratives will drive cortical activity similarly across participants, which we measured as inter-subject correlation of scalp potentials^39^. The effect is strongest during attentive listening and predictive of the recall performance. This suggests that cognitive processing of the narratives drives cortical activity. We hypothesized that this cognitive processing has a top-down effect on the autonomic system, affecting not only heart rate^35^ but also eye movements and pupil size (Hypothesis H1, solid arrows in Fig. 1C). We therefore predicted that these peripheral signals will be correlated to the electro-encephalographic scalp potentials (EEG) and be driven by cognitive processing of the stimuli. We will test this with signals recorded during the narrative, in both attentive and distracted conditions as well as during rest. Lastly we hypothesized a bottom-up effect of autonomic physiology on the brain (Hypothesis H2, dashed arrows in Fig. 1C). We will test this by correlating peripheral signals during conditions controlling luminance, saccades and breathing in separate control conditions.

A total of 67 participants were analyzed on data collected in two experiments (47 females, age 18-36 mean=22.49, standard deviation SD=3.71). The first cohort (N=29) looked at a blank gray screen during the narratives and during rest (Fig. 1B). The second cohort (N=38) was asked instead to look at a fixation cross with luminance matching the gray screen. All results reported here reproduced in these two cohorts, except where differences are highlighted.

**Figure 1:**
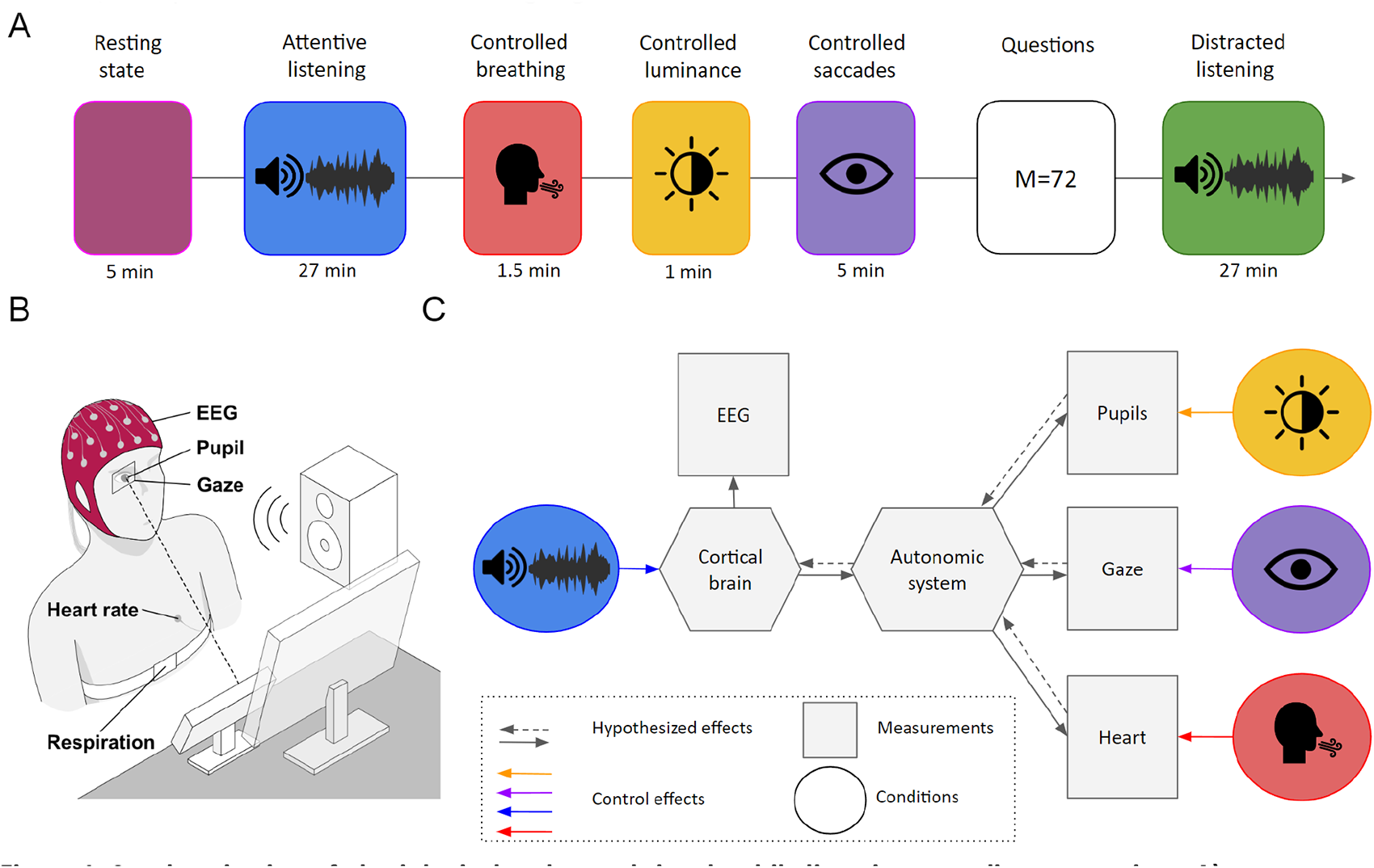
Synchronization of physiological and neural signals while listening to auditory narratives. **A)** Experimental paradigm: An initial 5 minutes of resting is followed by attentive listening to audio narratives (10 stories of 27 min duration), controlled breathing for 90 sec, controlled luminance for 60 sec (screen fluctuating between black and white at f=0.1Hz), controlled saccades (dots appear sequentially at 9-point on a grid with varying durations). Then questions are asked about the stories (4-10 questions per story) and lastly distracted listening to the same stories (but now counting backwards in decrements of 7 silently in their heads). **B)** Signal modalities collected: Scalp potentials of electroencephalography (EEG), Eye tracking (horizontal, vertical), pupil size, electrocardiography (ECG) and Respiration. **C)** Diagram of measurements, controlled effects and hypothesized effects.

### Saccade initiation and blink onset is affected by cognitive processing of the auditory narrative

Hypothesis H1 predicts that eye movements will be driven by the auditory narrative. In an effort to emulate natural listening conditions, for the first cohort of subjects, we did not require anything from the subjects other than to look at a blank gray screen during the listening task.

If the cognitive processing of the narrative elicits saccades we would expect them to be initiated at specific moments of the narrative. To test this we measured how similar saccade initiation was across listeners using intersubject correlation. We resolved this in frequency to determine at what timescale of the narrative this occurs. We find that saccade initiation did synchronize between subjects, but only during attentive listening (Fig. 2A). The time scale of synchronization of 0.1Hz or less is much slower than the average saccade rates of approximately 2Hz (Attentive 1.79 ± 0.96Hz, Distracted 1.91 ± 0.78Hz). This suggests that specific moments of the narrative tended to elicit saccades in multiple subjects and importantly, this is modulated by attention suggesting that this is not an automatic reflex to sound volume fluctuations.

To determine whether semantics of the narratives drive saccades, we follow the approach that has previously been used for scalp potential during continuous narratives^41, 42^. Specifically, we estimate a “temporal response function” (TRF) while participants listen to the narratives. A TRF captures the impulse response from an “input” signal to an “output” signal (see Methods). Here the “input” signal indicates word onsets (as a pulse train) and the “output” signal indicates the saccades (set to 1 during saccades and 0 otherwise). We find a significant response to words in the TRF at 100ms to 400ms when listeners are paying attention (Fig. 2B), however this is no longer the case when they are distracted, suggesting that processing words is a prerequisite for this increase of saccade likelihood. However, the number and the size of saccades during the narratives were not consistently different between the attentive and distracted conditions (Fig S2B and S2C, see also Section S2). We also tested if saccade rate fluctuations entrained to the auditory narratives as we had previously observed for video^51^. However, we do not find that the saccade rate changes in power (Fig. S2E) or in synchronization between subjects (Fig. S2F). This suggests that the basic statistics of saccades are largely unaffected by the narrative, but that subjects tend to saccade at specific moments of narrative when they are attending to the words.

Previous reports have linked blinking to pauses in continuous speech^52^. Do we see this effect here as well and does it depend on cognitive processing of the narrative? To test this we compute the ISC of blinking (Fig. 2C). We find a similar behavior to saccades, namely, blinking is locked to the narrative at a slow timescale of 0.05Hz-0.1Hz (Fig 2C). This is slower than typical blink rate (0.78Hz, see Fig. S3D), and the average word rate of the stories (2.90 ± 0.28 words/sec, see Fig. S6E), so blinks may occasionally be evoked at specific moments in the narrative and could be at the timescale of sentences (0.3 ± 0.29 sentences/sec, see Fig. S6F). As with saccades, we measured the TRF of blinks in response to word onset and found a significant response at 100 to 300 ms during the attentive condition only (Fig 2D). We conclude that both saccade initiation and blink onset in the absence of a visual stimulus, are likely driven by similar processes that drive scalp potentials during continuous speech ^41–43^.

**Figure 2:**
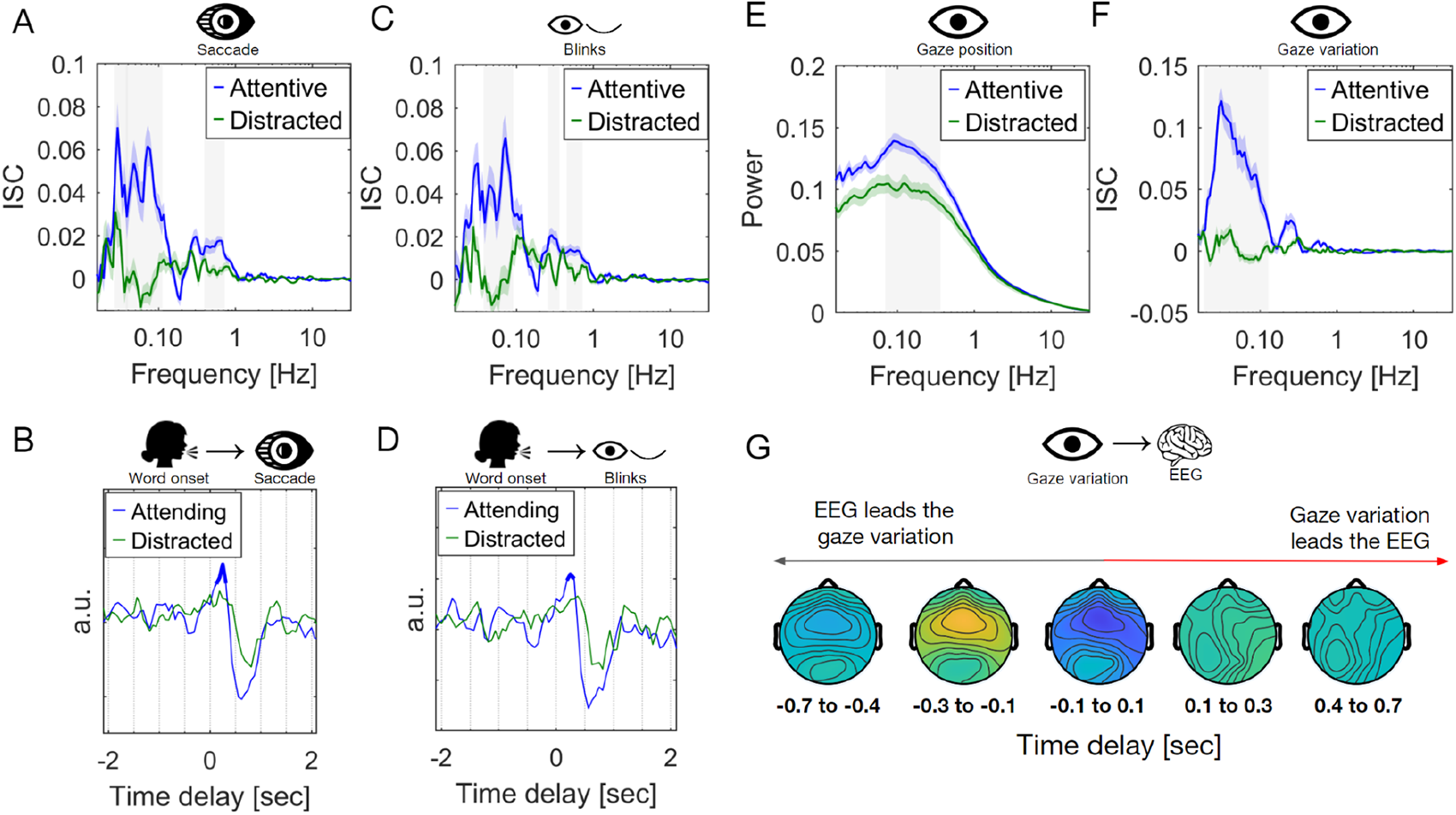
Eye movements during listening to auditory narratives. **A)** Frequency-resolved inter-subject correlation computed by first band-pass filtering the saccade signals (set to 1 during a saccade occurs and 0 otherwise) at different frequencies and then computing ISC averaged over stories and subjects. Colored shading indicates standard error around the mean (SEM) across N=29 subjects. Significance difference in ISC between the Attentive and Distracted condition is indicated as gray shading. Significance is established in each band using t-test, corrected for multiple comparisons using one-dimensional cluster statistics (p<0.01, N=29). **B)** Temporal Response Function using word onset (pulse train) as input signal and saccades as output signal (the saccade signal as panel A is used. **C)** Frequency resolved ISC of blinks computed between blink signals (set to 1 during a blink and 0 otherwise). **D)** Temporal Response Function using word onset as input signal and blinks as output signal. **E)** Power spectrum of gaze position computed by first band-pass filtering the gaze position signal, then computing the standard deviation for each subject and taking the mean over stories and vertical/horizontal direction and lastly normalized by the total power per subject across conditions. Significance difference between the Attentive and Distracted condition is established in each band using Wilcoxon rank sum test, corrected for multiple comparisons using one-dimensional cluster statistics (light gray area indicates p<0.01, cluster corrected). **F)** Frequency resolved ISC of gaze variation. Gaze variation was computed as the absolute value of the hilbert transformed gaze position after band-pass filtering. **G)** Temporal response functions of scalp potentials measured using each EEG electrode as output signal and the bandpass filtered gaze variation signal (0.1 to 1Hz) as input signal during the auditory narratives. Color indicates scalp potential fluctuations in response to fluctuations of pupil size (yellow and blue are positive and negative correlations). The dotted line indicates zero lag between the pupil size and scalp potentials. All data here is for Experiment 1, N=29 subjects.

### Gaze variation is modulated by attention and shows anterior-central neural activity

We also analyzed how gaze position fluctuated over time. A power spectrum of the gaze position signal reveals the amplitude of these fluctuations at different times scales (Fig. 2E) – we refer to these as gaze variations in the following. Attentive listening resulted in increased gaze variation around 0.1Hz (10 second cycles, see example video of these slow gaze variations <u>here</u>) as compared to hearing the narrative in the distracted condition. This is substantially slower than the speed of saccades or their rate of occurrence. These slow variations in gaze position suggest that subjects do not saccade more frequently, but simply that gaze position covers a wider field of view when attentively listening to the story.

After these findings on gaze variation, we repeated the experiment on a second cohort of participants (Experiment 2, N=38), but now we asked listeners to look at a fixation cross (with luminance equal to the background). Surprisingly, these slow gaze variations were still larger during the attentive versus the distracted listening conditions (Fig. S5C). Only during rest, where the only task was to maintain fixation, did we have the expected drop in gaze variation (Fig. S5A-B). Apparently, it is difficult for participants to focus on the fixation point when their attentional focus is on the narrative or the counting task.

To determine if these gaze variations are driven by the narrative, we measured their correlation across subjects during the narrative (Fig. 2F). During attentive listening, we see significant ISC in the range of 0.05 – 0.1Hz, and this is absent during the distracted conditions. This effect reproduces when subjects were asked to maintain fixation (Fig. S5D). This indicates that the moments of high and low gaze variation coincided across participants, but only if they are attending to the narrative. To measure the brain-body interaction between these gaze variations we computed the TRF predicting scalp potentials from the gaze variation signal (Fig. 2G). We find a positive anterior-central neural activity preceding fluctuations in gaze variation (100-300 ms). This differs substantially from the scalp potentials associated with gaze position (vertical Fig. S4B and horizontal Fig. S4C), blinks (Fig. S4D) and saccades (Fig. S4E). Here scalp potentials are largely following the onset of these movements, which reflects well known eye movement artifacts. In contrast, the anterior-central activity is predictive of subsequent changes in gaze variation suggesting a top-down effect of the processing of the narrative on gaze variation.

### Pupil size is affected by cognitive processing of the auditory narrative

As part of hypothesis H1 we also predicted a top-down effect of cortical processing of the narrative on pupil size. To test this, we measured TRF of scalp potentials in response to pupil size fluctuations for each electrode during attentive listening to the narratives. We see a clear “response”^1^ at positive and negative time lags (Fig. 3A), indicating that pupil fluctuations both precedes and follow neural activity with no clear causal direction. The spatial distribution of this correlated brain activity has at least two distinct patterns on the scalp: An earlier anterior-central negativity (-300 to -100ms) which is repeated positively (100 to 300ms), and a later bilateral occipital activity (400 to 700ms).

To determine which time-scale of fluctuations dominate this effect, we analyze the correlation between scalp potentials and pupil size, but now resolved by frequency for any time delay (Fig. 3B & 3D). We refer to this coherence measure as within-subject correlation (WSC)^36^. Correlation is evident in a broad frequency range from 0.3Hz to 3Hz. The associated spatial patterns indicate that the faster activity has an anterior-central distribution (Fig. 3B). Interestingly, when comparing the attentive vs distracted listening conditions we see no difference in the strength of WSC. This suggests that the pupil-cortex coupling is largely unrelated to cognitive processing of the narrative and instead, appears to be an endogenous link independent of an auditory stimulus.

The question remains, however, whether cognition drives autonomic activity reflected in the pupil. We test this by measuring inter-subject correlation (ISC) of the pupil, and resolve this by frequency (Fig. 3C). Significant ISC is present around the 0.1-0.5Hz band while participants attentively listen to the narrative, but not when they are distracted. Knowing that these narratives also entrain scalp potentials^39^, this suggests that the pupil is driven by cognitive processing of the narrative, via an endogenous link of cortical activity acting on the pupil. Consistent with this, we find that the overall strength of ISC of pupil size of an individual subject with the group is predictive of how well that individual recalls information about the narrative in the subsequent questionnaire (Fig. S2I). What is it in the narratives that could elicit pupil fluctuations?

As we did with saccades and blinks, we measured the response of pupil size to word onset using the same TRF formalis as before. We see a pupil dilation at a latency of 2s (Fig. 3E) and this effect is reduced when the participants are distracted from the narrative. Could this be associated with sound volume fluctuations which are known to elicit correlated fluctuations in scalp potentials.^43^ To test this we look at how the pupil size responds to sound volume fluctuations of speech. We find that the temporal response function of the pupil size shows non-zero responses at 1s latencies, but this is true for both attended and unattended speech (Fig. S6A-B). Thus, the response of the pupil to words implies that this is not just a low-level response to sound.

Another aspect that could drive pupil fluctuations is the complexity of the speech and associated listening effort^13^. As a measure of lexical difficulty we computed the word rate (number of words per second) and correlated that with the pupil size. We found a positive correlation in the frequency range of 0.03-0.4Hz (Fig. S6F-G) and a peak in the TRF around 1-1.5 seconds (Fig S6C-D), which are on the slower side of word rate fluctuations (Fig. S5E). Both these findings are consistent with our interpretation that cognitive processing of the speech drives the pupil via an endogenous link to cortical activity.

In conclusion these results indicate that the speech tracking phenomenon observed for scalp potentials^42, 56^ carries over to the pupil with both low-level and high-level features of the speech driving pupil size and these are modulated by top-down processes.

One caveat of our results on pupil size is that saccades and eye blinks are known to cause pupil size fluctuations^45, 46^. Indeed, we confirm the effect of saccade and blinks on pupil size (Fig. S1). Therefore, in the preceding analysis we removed any linear effect due to saccades and blinks from the pupil size measurements (see Fig. S1). Therefore, none of the effects in Fig. 3 can be attributed to the saccade and pupil effect reported in Fig. 2, except for the ISC effect which can capture nonlinear pupil “responses” that are similar across subjects.

**Figure 3:**
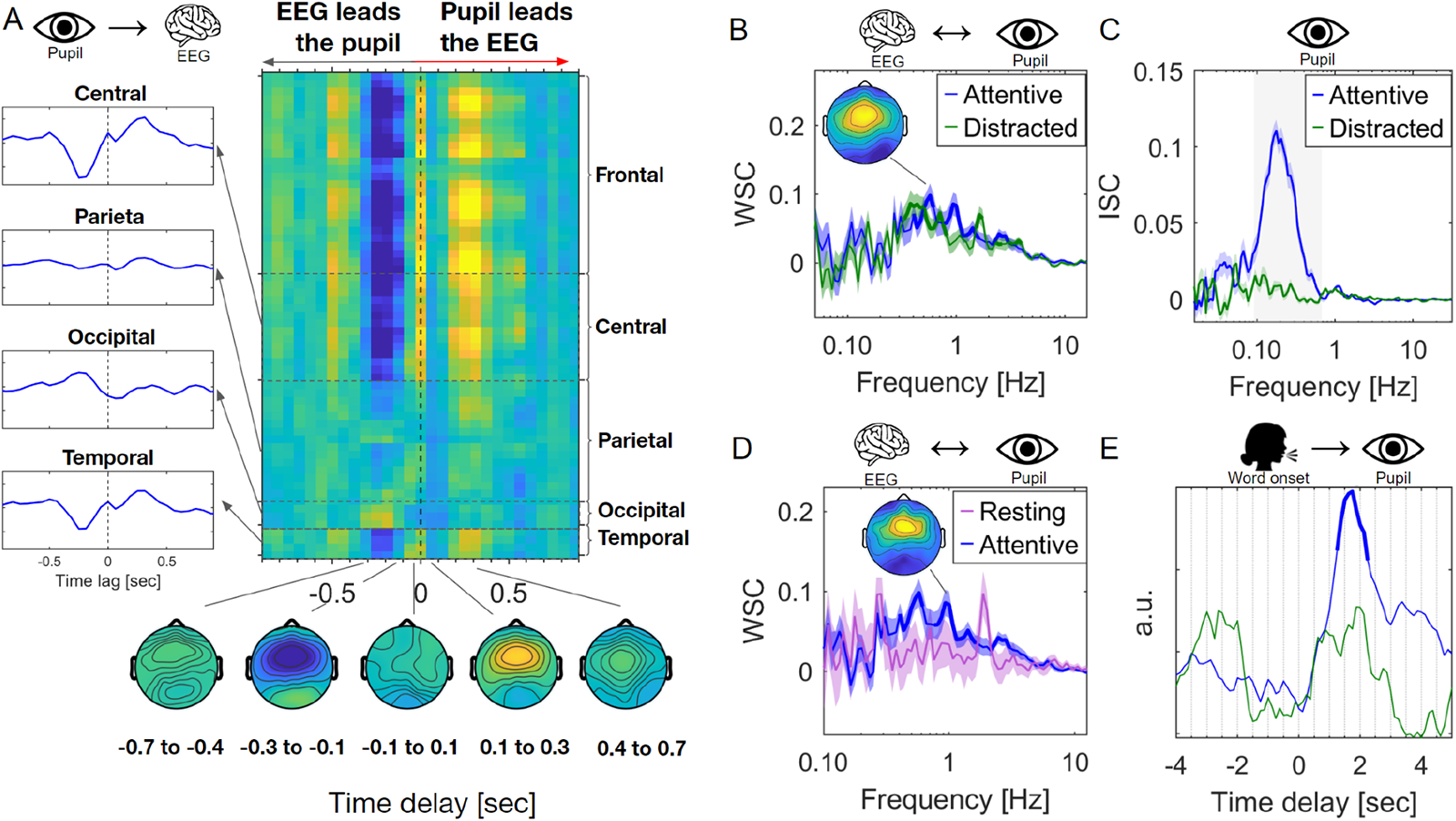
Pupil size during listening to auditory narrative: **A)** Temporal Response Functions using each EEG electrode as output signal and pupil size as input signal during the auditory narratives. Color indicates scalp potential fluctuations associated with pupil size (yellow and blue are positive and negative correlations). The dotted line indicates zero lag between the pupil size and scalp potentials (Experiment 1, N=29 subjects). The left panels are the average TRFs for the, central, parietal, occipital and temporal brain areas. **B)** Within-subject correlation between pupil size and scalp potentials during attentive and distracted listening to the narrative. Bold lines indicate significant WSC computed using t-test across subjects with cluster correction (p<0.01, N=29 subjects). **C)** Frequency-resolved inter-subject correlation computed by first band-pass filtering the pupil size at different frequencies and then computing ISC averaged over stories and subjects. Shading indicates standard error around the mean (SEM) across N=29 subjects. Significance of ISC between the Attentive and Distracted condition is established in each band using the same method as panel B. **D)** Within-subject correlation between pupil size and scalp potentials during attentive listening and resting state. Gray area indicates a significant difference between the Resting and Attentive conditions computed using t-test with cluster correction (p<0.01, N=29 subjects). **E)** Temporal Response functions using pupil size as output signal and word onset as input. Bold line indicates a significant difference in TRF between conditions computed using shuffle statistics with FDR correction (p<0.01, N=29 subjects).

### Heart rate is affected by cognitive processing of the auditory narrative, and correlates with scalp potentials

Hypothesis H1 of a top-down effect of cognition on autonomic function also predicted a correlation of scalp potentials with the heart rate signal. Instantaneous heart rate can be computed as the inverse of the beat-to-beat interval (see Methods). We again measure TRFs, now with the instantaneous heart rate as “input” and scalp potentials as “output”. We find non-zero responses to heart rate, mostly leading the neural signal (Fig 4A). Interestingly, we see again a anterior-central component in the scalp potentials that is anticipated by the heart rate by almost 1s. The same pattern is present for responses in the distracted and rest conditions, suggesting that this heart-cortex link is endogenous. The similarity of the frontocentral component for pupil and heart is suggestive that there is a common source in the brain for this effect (compare Fig. 3A and 4A).

As predicted by hypothesis H1 we also find that the narrative synchronizes heart rate between subjects (Fig 4B), but only during the attentive listening condition. This replicates our previous findings on video^35^ and confirms that it is the cognitive processing of the narrative that drives these slow-frequency (0.1Hz) heart rate fluctuations. Incidentally, we additionally recorded electrodermal activity, which is another peripheral signal affected by sympathetic drive^57^. As with other peripheral signals, we find that it synchronizes between subjects during the narrative below 0.1Hz, but only while attentively listening (Fig. S7A-B).

So far we have seen cognitive effects on heart rate, however recent work has also shown cognitive and perceptual effects on individual beats and their association with cortical activity^58^. The latter have been measured as Heartbeat Evoked Potentials (HEP)^59, 60^. We find weak but significant differences of the HEP bilaterally over posterior cortex when comparing attentive listening with the rest condition (Fig. S9A), but we did not find evidence of attentional modulation of the HEP (Fig. S9B).

A common measure of the effect of cognition on heart rate is heart rate variability, which, when resolved by frequency, is simply the power spectrum of the instantaneous heart rate (Fig. 4C). The well-established high (HF-HRV) and low frequency (LF-HRV) components of heart rate variability^61^ are quite evident here, but we see no effect of attention. We note that the high-frequency component corresponds to rhythmic breathing which modulates heart rate at the breathing frequency^62^. In our data this is most evident in the within-subject correlation between respiration and heart rate (Fig. 4D). We see the respiratory sinus arrhythmia as a peak at 0.3Hz. However, the synchronization of heart rate across subjects is not the result of synchronized breathing, as there is no reliable ISC for breathing during the narratives (Fig. S7C-D). WSC between respiration and heart rate is also not modulated of attention (Fig. 4D). This is also evident in the instantaneous heart rate signal aligned to peak inhalation (Fig. 4E).

Respiratory volume has recently been linked to arousal and it has been shown to correlate with the global signal in fMRI at ultra-slow fluctuations^63^. In our data we find a synchronization of respiration volume across subjects, with a modulation by attention at 0.04Hz (Fig. 4F and S7F). However we do not find any coupling between scalp potentials and respiratory volume at these very slow fluctuations (Fig. S7G), but we can not rule out limitations of the EEG equipment in this low frequency range.

**Figure 4:**
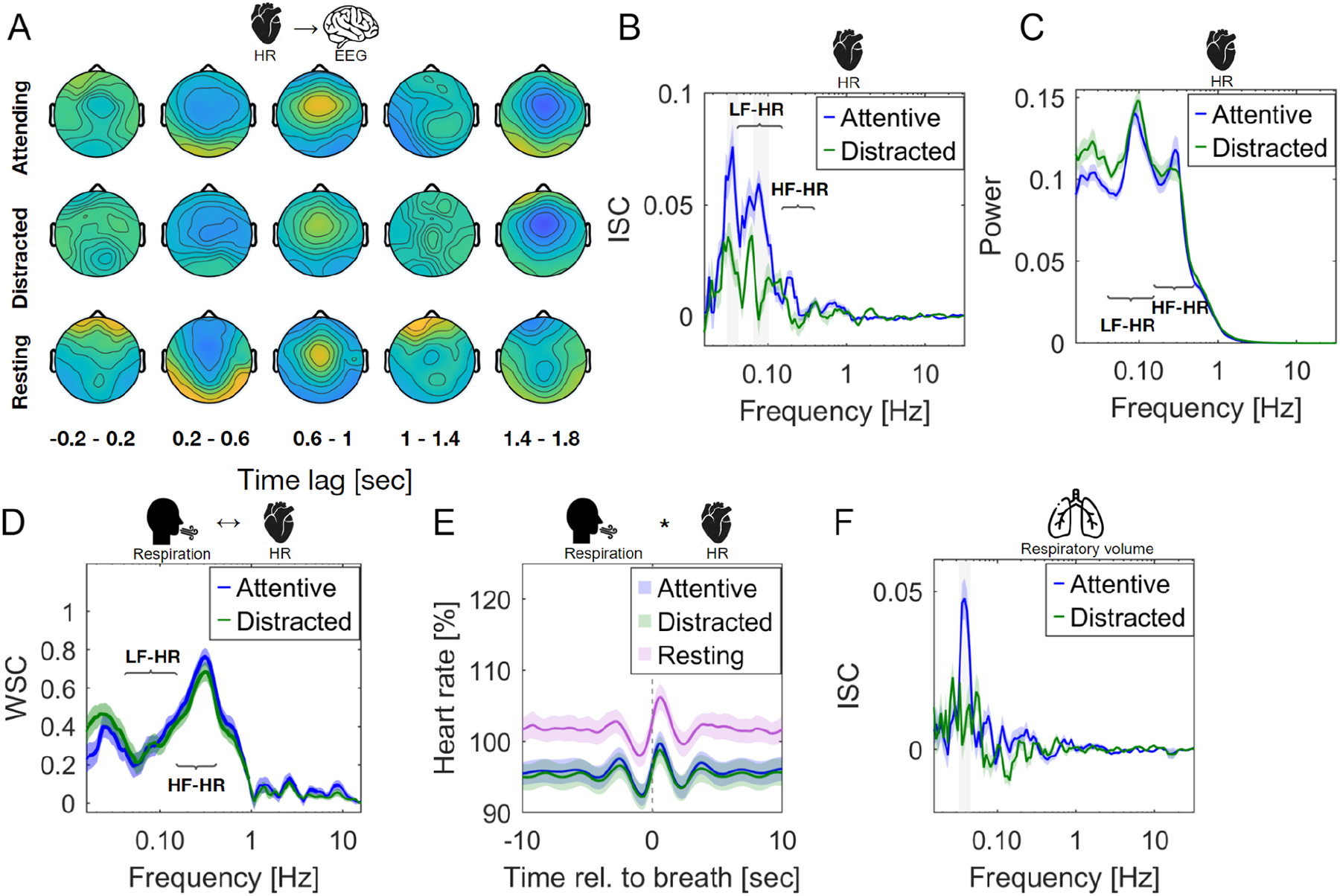
Heart rate and breathing during listening to auditory narratives. **A)** Topographical plot of temporal response function between heart rate and scalp potentials, similar to Fig. 2A. Positive lag indicates that heart rate fluctuations lead scalp potential fluctuations (N=29). **B)** Frequency-resolved ISC of heart rate. Colored shaded area indicates SEM across N=29 participants. Gray shaded area indicates frequency bands with significant differences between attend and distracted listening conditions. Significance of ISC is established in each band using t-test, corrected for multiple comparisons using one-dimensional cluster statistics (p<0.01, N=29). LF-HR indicates the Low Frequency heart rate region, HF-HR is the high frequency heart rate range. **C)** Power spectrum of heart rate, computed as the normalized power (dividing the power in each frequency bin with the aggregate power of the Attentive and distracted condition). Significance between the Attentive and distracted condition was compared as Panel B, however using the Wilcoxon rank sum test instead of the t-test) **D)** Within-subject correlation between respiration and heart rate. (statistics and error bars computed like panel C. **E)** Breathing-evoked heart rate fluctuations relative to baseline (10 seconds average before each story was played) (reverse correlation). **F)** Synchronization spectrum for respiratory volume. Error bars and statistics were computed in a similar fashion as panel B.

### Bottom-up effect of heart rate control, but not pupil control

Hypothesis H2 posits a bottom-up effect of autonomic function. By this we mean that brain nuclei controlling, e.g. pupil size, may also affect heart rate, or vice versa, in the absence of a top-down cognitive drive. To test this, we analyzed data collected during controlled breathing and controlled luminance, with both conditions in silence and compared these to resting state. As the controlled interventions were time-aligned across subjects (see Methods), we can again use inter-subject correlation to capture reliable responses to these interventions. Assuming that effects are the same across subjects this approach can capture linear as well as non-linear or delayed responses to the interventions.

In the controlled luminance condition the brightness of the screen was modulated at a rate of 0.1Hz. This elicited an obvious synchronization of pupil size across subjects (Fig 5A), as well as a synchronization of eye movements (gaze variation and saccade rate). Note that there was no fixation cross, so it is conceivable the change in brightness may have prompted subjects to move their gaze in a predictable way, and synchronization in the EEG is possibly just the result of eye movement artifacts.

### Importantly, we find no effects on heart rate or breathing

Synchronized breathing on the other hand had broad-ranging effects, synchronizing not only heart rate but also pupil size, saccade rate and EEG (Fig. 5B). We can directly compare the bottom-up effect of controlled breathing and luminance by computing the WSC between heart rate and pupil size (Fig. 5D and Fig. 5E) and compare that to when participants are at rest. Here we see a significant peak in the WSC at 0.1Hz of the controlled breathing. This is the same frequency participants were asked to breathe. However, this correlation is not present for the controlled luminance condition (with luminance modulated at the same frequency). This suggests that while brain nuclei controlling pupil size have a fairly narrow effect, the effect of modulating heart rate using respiration seems to spill over to a number of autonomic signals. Consistent with this, we find that the correlation of heart rate with pupil size remains when participants were at rest with a peak at 0.3Hz (Fig. 5E), corresponding to the natural breathing rhythm.

When we listen to auditory narratives, and breathe at our natural breathing rate, the pupil heart rate link remains (Fig. 5F). However, we do not see any significant effect of attention, suggesting an endogenous link. We also analyzed the relative time delay between these coherent fluctuations using the formalism of TRF. In this case we model the linear response of the pupil size onto heart rate and vice versa (Fig. 5C). In both instances, we obtain non-zero responses with fluctuations in heart rate that are ahead of fluctuations in pupil size. Taken together these results suggest that the autonomic control of heart rate has a bottom up effect on other autonomic signals confirming H2, but the same is not true for pupil control.

**Figure 5:**
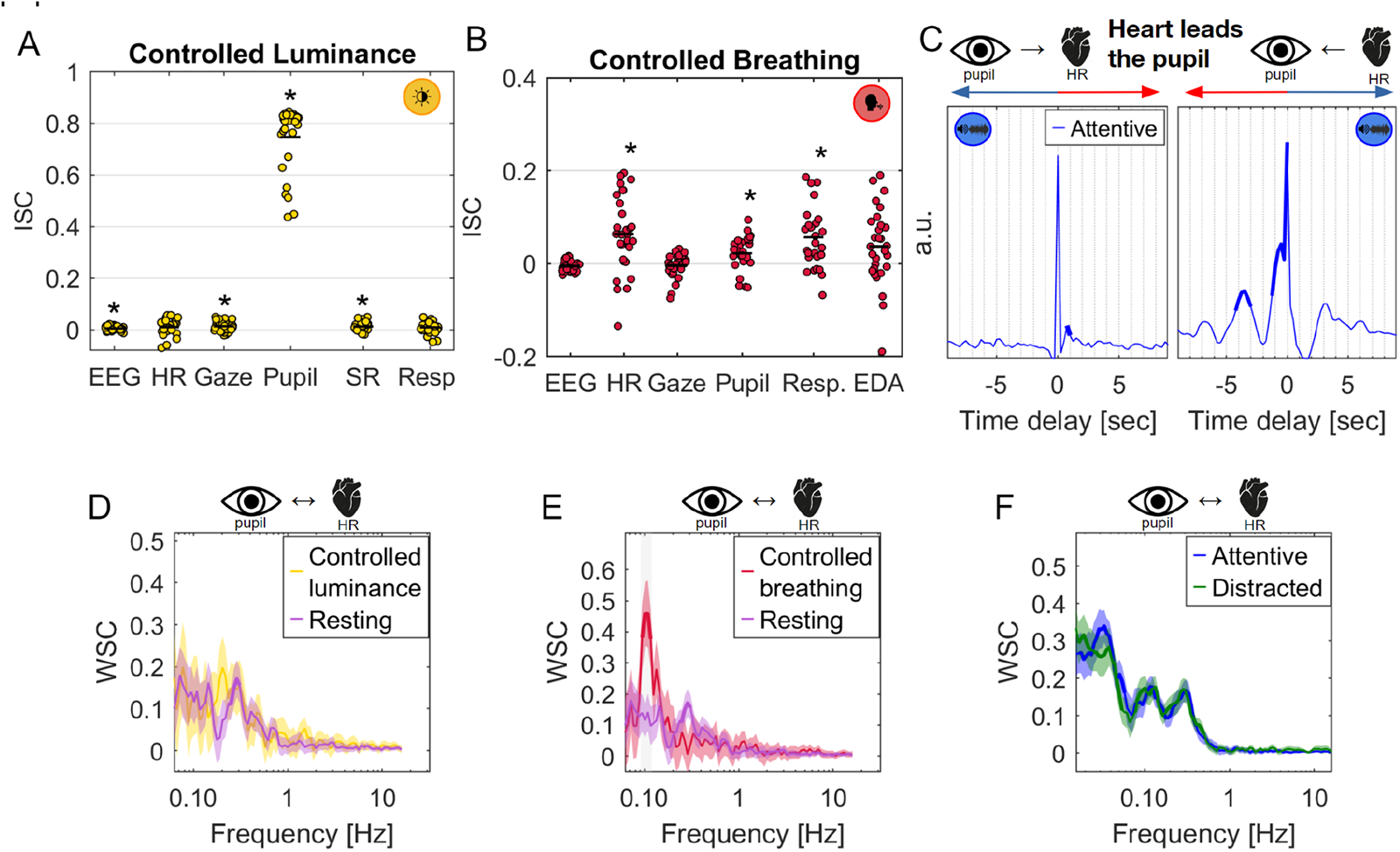
Cross-modal effects of controlled luminance and controlled breathing. **A)** Inter-subject correlation during the controlled luminance condition. ‘*’ indicates the ISC values are significantly different from zero using a t-test (p<0.01, N=29) **B)** Inter-subject correlation during the controlled breathing condition. **C)** Temporal response function between heart rate and pupil size during attentive listening. Left panel models the linear response of pupil size (as output) to heart rate (as input). The right panel models the reverse. Non-zero response (for positive lags in the left panel, and for negative labs in the right panel, red arrow) indicates that fluctuations of heart rate are ahead of fluctuations in pupil size. **D)** Within-subject correlation between pupil size and heart rate during controlled luminance condition and resting state. **E)** same as panel D but for controlled breathing condition compared to resting state. **F)** same as panel E but for attentive and distracted listening conditions.

### Bottom-up effect of eye movements on heart rate

The other peripheral signal we have explored was eye movements. A bottom-up effect following hypothesis H2 might predict that eye movement has an effect on heart rate. To test this we asked subjects to follow dots on the screen that jumped in position every 2 seconds. Measuring ISC shows entrainment for a number of signals (Fig. 6A). Eye movements obviously synchronize saccade rate and pupil size, but also entrain EEG, presumably due to eye movement artifacts on the scalp potentials. However, the significant ISC we observe for heart rate can not be trivially explained. When resolved by frequency, we see a clear synchronization at the 0.5Hz rate of saccades (Fig. 6C), scalp potentials (Fig. 6B), pupil size (Fig. 6D) and lastly heart rate (Fig. 6E), suggesting a bottom-up effect of saccades on individual heart beats. A 0.5Hz oscillation in heart rate reflects a slow beat followed by a fast beat, in synchrony with the rhythmic saccades. In the second experiment we repeated this intervention with different rhythms of saccades (0.2 Hz, 0.5Hz, 1Hz and 2Hz), but only found significant synchronization for 0.5Hz (Fig. S8A-D). This suggests that saccades have an instantaneous effect on individual heart beats, that manifests most clearly when saccades follow the natural rhythm of the heart (Fig. 4C).

**Figure 6:**
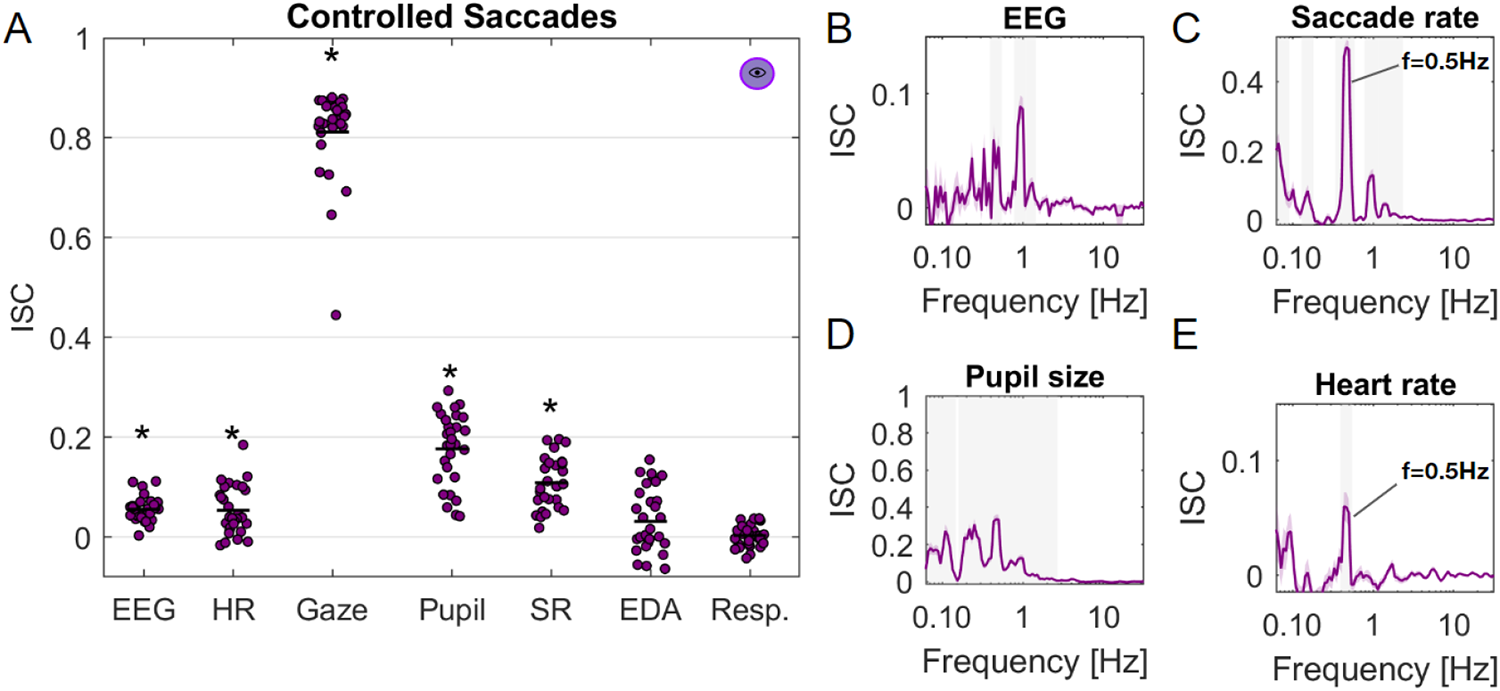
Rhythmic saccades modulates heart rate. **A)** Intersubject correlation of various modalities during the controlled saccade experiment. Participants followed a dot jumping on the screen in a 9-point grid every 2 seconds. ‘*’ indicates the ISC values are significantly different from zero using a t-test (p<0.01, N=29). **B)** Frequency-resolved ISC for EEG during the controlled saccade experiment. Line indicates the average across N=29 participants, the color-shaded area indicates the standard error of the mean. Significance of ISC was computed using a t-test in each frequency band, corrected for multiple comparisons using one-dimensional cluster statistics (Gray shaded area indicates p<0.01, cluster corrected). **C)** same as panel B now for saccade rate during the controlled saccade rate experiment. **D)** same as panel B now for pupil size during the controlled saccade rate experiment. **E)** same as panel B now for heart rate during the controlled saccade rate experiment.

## Discussion

We set out to test whether pupil size and eye movements are modulated by listening to an auditory narrative. We had seen this previously for viewing of audiovisual narratives^36^ but it was not obvious that it should replicate in the absence of a visual stimulus. We hypothesized that these signals are peripheral markers of arousal, and as such, should be similarly driven by top-down effects of cognition. We confirmed this here for pupil size and heart rate, and for saccade initiation and blinks.

Surprisingly, we found slow variations of gaze position on a time scale of 10s, with an amplitude that is modulated by the narrative: not only did attentive listening increase gaze variation, but the specific moments of high and low variation during a narrative were similar across listeners, suggesting that the semantic content of the narrative affected gaze variation. We draw the same conclusions for fluctuations of pupil size and heart rate: inter-subject correlation at a similar time scale of 10s was modulated by attention, and it was predictive of memory of the auditory narrative. Timing of blinks and saccades also entrained to the narrative at similarly slow timescales. Together, these results suggest that cognitive processing of the narrative drives autonomic function, perhaps reflecting varying levels of arousal. The link between cognition and autonomic arousal is consistent with the notion of allostasis, which is the efficient regulation required to prepare the body for future demands.^64^ In this view, cognitive processing of the narrative may be driving autonomic function to prepare the body to act.

We also set out to determine whether there is a bottom-up effect, whereby a low-level drive of these peripheral signals has an effect on the other autonomic signals. This is by no means a trivial proposition. While there is some overlap in brain nuclei known to control pupil size, heart rate and eye movements, the control networks for the corresponding effector muscles are quite distinct, and bottom-up afferent connections are not as clearly established. Breathing has a well-established effect on heart rate^62^, and is subject to a combination of central control and peripheral effect such as the baroreflex^23^. But based on the neuroanatomical literature alone it was not obvious that breathing would affect pupil size, yet this is what we found here. Another well-established, low-level effect is pupillary light reflex^1^. Luminance fluctuations might have affected arousal and thus driven heart rate. We did not see this here. Instead we found that heart rate fluctuations preceded pupil fluctuations, and in fact, they preceded slow cortical potential fluctuations. This suggests that cardiac control has a broad bottom-up effect on arousal, which pupil control does not. Indeed, there is a long list of studies arguing for a bottom-up effect of cardiac function of the brain^29, 30, 65^, with the effect often attributed to arousal^66, 67^.

Our results linking heart rate fluctuations to pupil size are consistent with previous studies that show a transient increase in pupil size and heart rate following a behavioral response^68^. Additionally, conscious perception of surprising auditory stimulus leads to both an immediate advanced heart beat^26^ and somewhat slower pupil dilation effect^16^. On this short time scale of individual heart beats, there also appears to be a modulation perceptual performance and evoked potentials with the time scale of a single cardiac cycle^58^. We do find a small enhancement of heartbeat evoked potentials during attentive listening as compared to the resting state. However the heartbeat evoked potential was not modulated by the attentional state during the narrative, as we had found with the ISC of heart rate. Thus, it may have resulted from the mere presence of sound.

We found that scalp potentials correlate with heart rate, pupil size and gaze variation. The spatial distribution for all three has a anterior-central distribution, suggesting a common cortical source. The anterior-central distribution is reminiscent of the “error related negativity” and is consistent with a source in anterior cingulate (ACC)^69^. In a recent integrative theory of the locus coeruleus-norepinephrine function they argue that that ACC plays a role in optimizing the utility between short timescale LC activity (phasic pupillary responses) and long timescales (tonic pupillary responses) which are associated with disengagement from task-demand. The slow fluctuations we find synchronizing between people could be related to these switches between internal and external attention. This interpretation is consistent with the finding that inter-subject correlation of scalp potentials during audiovisual narratives correlates positively with fMRI activation of ACC, among other brain areas^70^. Inter-subject correlation of fMRI activity in ACC is also predictive of performance in a quiz on the material presented in educational videos^71^, similar to what we found previously with EEG^72^. The power of heart rate fluctuations in the low-frequency band (LF-HRV), which is often associated with sympathetic arousal^61^, correlates positively with fMRI activity in ACC^73^, and HR itself correlates with fMRI in midline brain structures including the ACC^27^. Indeed the ACC is thought to be one of the cortical areas regulating autonomic function^29^. Additionally, activity in the ACC is modulated by activity in locus coeruleus^74^, which is tightly coupled with pupil size and often taken as an indicator of central arousal^5^. Activity in the ACC is also associated with conflict monitoring, including when just perceiving conflicting stimuli passively^69^. Here we found that words that are semantically dissimilar to the preceding words elicit pupil dilation, similar to the behavior of scalp potentials associated with semantic conflict (albeit with a broader central distribution)^42^. Taken together, the present results and prior literature suggests sympathetic arousal as result of detecting conflict, or even just unexpected semantics in the narrative. But given the broader involvement of the ACC in cognitive function, and its central role in the “somato-cognitive action network”,^75^ this may just be one of several aspects in the narrative that drove these peripheral autonomic signals.

As an aside, we want to highlight that in addition to semantic dissimilarity, we found that fluctuations in word rate and sound amplitude correlate with fluctuations in pupil size, and that this depends on attention. Such effects have been extensively reported for EEG^40^ and they replicated here for pupil size. This may open up opportunities for new research on speech processing^41^ and auditory attention decoding^76^ using the pupil in addition, or instead of EEG.

In analyzing eye movements we were motivated by the recent proposal that saccade initiation and pupil dilation are subject to a common central drive^15^. Saccades are driven by activity in superior colliculus, which receives input from the locus coeruleus, and that in turn is known to affect heart rate^4^. Given these overlapping brain nuclei, there was a possibility that cognitive processing of the narrative might have a top-down effect on saccade initiation. Conversely, the overlapping control nuclei may mediate a bottom-up effect of saccades on heart rate, or heart rate on saccades, irrespective of cognition. Indeed, we found a top-down effect of cognition on saccade initiation. Our observation that saccades and blinks entrain to the narrative during attentive listening is consistent with the recent reports that gaze position, saccades and blinks correlated with low-level features of continuous speech.^77–79^ We also found a bottom-up effect of saccades on individual heart beats. This is consistent with a previous observation that saccades tend to align in time with the cardiac cycle during visual search^80^. A link between saccades and heart rate has also been postulated in the context of “eye movement desensitization and reprocessing therapy”. This intervention asks patients to recall traumatic experiences while eliciting rhythmic eye movements (left/right movement approximately every 0.5 seconds). There are some reports that this intervention lowers heart rate during the procedure^81, 82^. We find that heart beats entrain to saccades only if they are repeated every 2 seconds, so the relationship to this clinical intervention is unclear.

An unexpected result is that gaze variation was larger during attentive listening and was driven by the narrative. However, neither saccade rate nor saccade size differed consistently with the attention manipulation. This suggests that low-level control of eye movements was not altered, but simply that our gaze canvases a wider area of the visual field when we naturally listen to a story. This result on gaze variation is consistent with a recent study, which shows that gaze variation (about a fixation cross) during pre-stimulus baseline correlated negatively with alpha power in the EEG^83^. Both alpha power and gaze variation were predictive of subsequent memory of the upcoming visual scene. Alpha is generally considered a measure of cortical inhibition.^84^ These results are therefore consistent with an interpretation of gaze variation as a measure of arousal. Eye movements are not required for the task of listening to a narrative, and therefore are incidental to the task. Previously we also found that viewing an audiovisual narrative entrains head movement amplitude, which would also seem to be incidental to the viewing task^36^. Movements unrelated to a task have recently been shown to dominate neural activity in cortical and subcortical brain areas in rodents^85, 86^. But the relationship of incidental behaviors to arousal has been unclear thus far. Here we have identified a clear association of a readily observable incidental behavior with traditional markers of autonomic arousal.

### We want to close the discussion with a few caveats

Our results on pupil size are similar to the results on saccade initiation and blinks. To rule out that this is a simple consequence of the known effects of saccades and blinks on pupil size^45, 46^, we removed linear correlations between these signals prior to analyzing pupil size. Of course it is possible that we did not adequately account for these interactions. On the other hand, these interactions are consistent with our hypothesis, and prior theory, that links saccade initiation and pupil to a common physiological drive related to arousal.^15^

It is important to acknowledge that the fluctuations driven by the narrative for both heart rate and pupil are relatively slow (∼10s) as compared to the link we find between these peripheral signals and the brain (∼1s). Interestingly, we do not detect pupil ISC in the faster 1Hz band, which we previously observed with audiovisual stimuli^36^, suggesting that these faster pupil fluctuations were due to the dynamic of the visual stimulus and/or saccades. An alternative explanation of our results at these slow time scales is that increased heart rate (and breathing) increases blood oxygenation. There is a global signal in fMRI at infra-slow timescales (<0.1Hz) that correlates with respiratory volume^87, 88^ and respiration volume/rate correlates with neural activity^89^. However, these slow fluctuations in fMRI signals that are attributed to fluctuations in arousal are in the infra-slow times-scales 0.01-0.1Hz^63^. We did find entrainment in respiration volume to the narrative at similarly low rates (0.04Hz) with a clear effect of attention. Breathing rhythms themselves (at 0.3Hz), which modulate heart rate^90^ were not entrained by the narrative, consistent with our previous results^35^. In total, while the breathing cycle itself had no effect, it is possible that heart rate is affected by breathing volume. Either way, respiration volume appears to be another signal reflecting the top-down effect of cognition on arousal similar to heart rate and gaze variation. However, respiration volume likely does not play a major role in the top-down effects we have found here on pupil size at a faster time scale (0.1Hz).

When measuring correlation of neural activity with the peripheral signals we have focused on fluctuations of the raw potentials and not their oscillatory power. The motivation for this is the extensive literature on these “evoked responses’’ during continuous natural stimuli, such as speech and videos^41, 54, 70, 91, 92^. We refer to them as evoked responses because they are the equivalent of event related potentials (ERP), except they are evoked by a continuous stimulus, rather than discrete events. Compared to conventional evoked responses (either to continuous or discrete stimuli) the time scale of fluctuations reported here are relatively slow. We ascribe this primarily to the relatively slow time scales of the peripheral signals we are correlating with. Nevertheless, the question arises as to what these slow fluctuations represent. The fluctuations are comparable to “slow scalp potentials”, which have been reported in the context of cognition and action in early EEG work^93^. Evoked responses are generally thought to reflect large scale waves of synaptic activity (as opposed to neuronal firing), but the physiological origins of slow fluctuations are not well understood. A more recent report suggests that fluctuations in the 0.1Hz range correspond to hemodynamic activity in the brain^94^, although it is worth noting that fluctuations we observed in the EEG are not as coherent as those reported there. Incidentally, from a practical point of view, and for future work, it is worth noting that not all EEG hardware will record at very low frequencies due to limitations of the electronic hardware.

The time scale may also coincide with electro-dermal activity, which is invoked in some of the literature on autonomic function and “stress”, with possible cognitive aspects^57^. Indeed, we found that electro-dermal activity (measured on the index and middle fingers of the non-dominant hand) synchronized during attentive listening to the narrative, but not any of the other conditions. Electro-dermal activity is due to fluctuations in skin resistance that manifests in slow fluctuations of voltages measured on the skin surface of the arm/hand/finger. However, if EDA was to affect scalp potentials, this signal would be removed during recording as the ground electrode is placed nearby and EDA is presumed to be uniform across the surface of the skin. Instead, the potentials we measure have diverse topography across the scalp, consistent with a direct cortical origin of this slow activity, and inconsistent with EDA.

By simultaneously recording brain activity and a variety of peripheral signals we investigated how the brain and body interact during natural behavior like story listening. In the past many of these phenomena have been studied in isolation within neuroscience, physiology and psychology. We believe that the present results add to growing evidence for an interaction between body and mind. But we also see a relevance for research on auditory processing, autonomic function, cognitive effects of arousal, and slow signal fluctuations observed in the brain.

## Acknowledgement

This work was funded by the National Science Foundation through grant number DRL-2201835. We want to thank Alexandria Hoang for her help with collecting some of this data. We would like to thank Vinay S. Raghavan for alerting us about the potential confound of blinks on pupil size.

## Author contributions

J.M. and L.C.P. contributed to the conception and design of the study. J.M. contributed to the acquisition of data. J.M. and L.C.P., contributed to the analysis of data. J.M. and L.C.P. contributed to the writing of the manuscript and/or the figures.

## Star Methods

### Key Resource table

| REAGENT or RESOURCE | SOURCE | IDENTIFIER |
| --- | --- | --- |
| **Deposited data** |  |  |
| Raw and analyzed data | This paper | Will be made available upon publication |
| **Software and algorithms** |  |  |
| Processing and analysis scripts | This paper | Will be made available upon publication |

### Resource availability

#### Lead contact

Further information and requests for resources should be directed to and will be fulfilled by the lead contact, Jens Madsen.

### Materials availability

This study did not generate unique reagents.

### Experimental model and subject details

All experiments were carried out at the City College of New York with the approval from the Institutional Review Boards of the City University of New York. Documented informed consent was obtained from all participants at the start of the experiment. Experiment 1 had N=32 participants (16 Female, age 19-36, M=23.69, SD=4.42; 3 subjects were removed due to bad timing, missing or bad signal quality). Experiment 2 had N=43 subjects participating (47 females, age 18-30, M=21.58, SD 2.81 years; 5 subjects were removed due to bad timing, missing or bad signal quality).

### Method details

#### Stimuli

The stories, which we used in a previous study^36^ were autobiographical narratives selected from the StoryCorps project aired on National Public Radio (NPR) and the New York Times Modern Love series (N=10, 2-5 min each, total duration of 27 min). During the presentation of the audio a gray background was presented on the screen for Experiment 1 and an isoluminant fixation cross was added at the center of the screen for Experiment 2. The stories were selected to be engaging and have an emotional narrative. 5 stories came from the New York Times’ Modern Love episodes: “Broken Heart Doctor”, “Don’t Let it Snow”, “Falling in Love at 71”, “Lost and Found”, and “The Matchmaker”, and 5 stories came from StoryCorps’ animated shorts: “Eyes on the Stars”, “John and Joe”, “Marking the Distance”, “Sundays at Rocco’s” (depicted in Fig. 1a), and “To R.P. Salazar with Love”).

#### Procedure

Participants listened to the stories seated comfortably in a sound-attenuated booth with white fabric walls and normal ambient LED lighting placed around the subject. The stories were played through stereo speakers placed at 60° angles from the subject next to the 27” monitor, all at a distance of approximately 60 cm from the subject. The procedure of the experiment can be seen on Figure 1.

##### Attentive and distracted listening

In the attentive condition participants were asked to attend to the story while looking at the screen (Experiment 1) or keep fixation on the isoluminant fixation cross centered on the screen (Experiment 2). In the distracted condition participants listened to the stories again but they were asked to count backwards silently in their mind in steps of 7, starting from a random prime number between 800 and 1000. This condition ensures that the stimuli and surroundings are identical, only changing the participants cognitive state.

##### Controlled breathing

To test the bottom-up effect of heart rate, we asked participants to breathe synchronously at a rate of 0.1Hz by showing them a waveform of the desired breathing pattern while looking at a centered fixation cross. To eliminate any visual input while they carried out the synchronized breathing task they were asked to continue breathing at the same rate while a uniform gray screen was presented. Data was analyzed during 60 seconds of this free breathing period (Experiment 1). In the second experiment the free breathing period was increased to 90 seconds and subjects were asked to look at a centered fixation cross (equal luminance to the gray background).

##### Controlled luminance

To test the bottom-up effect of pupil size, we asked participants to keep their gaze centered on the screen while we displayed a slowly flashing screen going from white to black (in 5s and back to white in another 5s, so a frequency of 0.1Hz) for a total of 60 seconds. Prior to the start of the flashing screen we displayed a uniformly gray screen for 60 seconds. To remove any transient effect of the pupillary response we did not analyze the initial 5 seconds of the condition.

##### Controlled saccades

To test the bottom-up effect of eye movements, participants were asked to direct their gaze to a sequence of dots shown on a screen one at a time in 4 different conditions. In each condition dots were either shown for 0.5, 1, 2 or 5 seconds aiming to produce a saccade rate of 2, 1, 0.5 or 0.2Hz respectively. For each condition a sequence of 9 dots was used (repeated multiple times), which was the same sequence across participants of randomly chosen dot positions, sampled from a 9-point grid (3 by 3 dot design with equidistant points on the screen). To ensure each condition lasted around 90 seconds the 4 different conditions (0.5, 1, 2 or 5 second) the sequence of 9 points was repeated 20, 10, 5 and 4 times respectively. To minimize any pupillary response due to luminance change when participants were saccading between dots, each dot was chosen to be isoluminant (equal luminance) to the gray background screen.

##### Questionnaire

After subjects listened to all stories in the attentive condition and carried out control conditions they were asked to answer questions about each story (8-12 questions per video, total 72 questions). The list of questions is available in a previous study^36^.

### Quantification and statistical analysis

#### Alignment of multimodal recordings

For segmentation of the physiological signals we used common onset and offset triggers, in addition, a beep sound was embedded right before and after each story which were recorded using a StimTracker (Cedrus) to ensure precise alignment across all subjects. To enable all modalities to be on the same timescale we used the lab streaming layer (LSL) protocol. In addition triggers were sent to both the eye tracking and ExG recording systems, by collecting the timestamps from each system when the triggers were sent, we can estimate a linear regression model converting timing of triggers from one modality to the other. This is specifically important when we use the EyeLink system as the quality of the data in “Streaming” mode could be diminished.

#### Recording and preprocessing of gaze position, gaze variation and pupil size

Gaze position and pupil size were recorded using the Eyelink 1000 eye tracker (SR Research Ltd. Ottawa, Canada) with a 35mm lens at a sampling frequency of 500 Hz. Subjects were instructed to sit still while the experiment was carried out, but were free to move their heads, to ensure comfort (no chin rest was used). A standard 9-point calibration scheme was used utilizing manual verification. Stable pupillary responses were ensured by adjusting the background color, dots used for calibration or interventions, or text in the instructions to have equal luminance (iso-luminant). Blinks and saccades were detected using the algorithm of the eye tracker. Artifacts, blinks and 100ms before and after were filled with linearly interpolated values. Effects of blinks and saccades on the pupil signal were removed using linear regression. To regress out the effects of the saccades and blinks on the pupil signal we first downsampled the pupil signal to 100 Hz, the Temporal Response Function was then estimated for saccade and blink offsets in the interval from -4 to 5 seconds independently (this is the region we have found the effects of saccades and blinks on the pupil size. The linear effects of these two TRFs were then regressed out simultaneously as blinks and saccades have interactions. We compute the gaze variation as follows 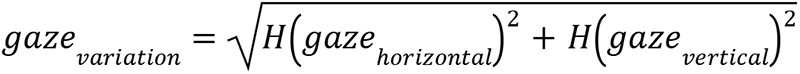, where *H* indicates the hilbert transform and *gaze* is the gaze position after band-pass filtering. Both the gaze position, gaze variation and pupil size were downsampled to 128 Hz.

#### Recording and preprocessing of EEG

The EEG was recorded at a sampling frequency of 2048 Hz using a BioSemi Active Two system. Participants were fitted with a standard, 64-electrode cap with electrode placements following the international 10/10 system, where the ground electrodes were located next to POz. To later remove any facial or eye movements artifacts from the EEG signal, we additionally recorded the electrooculogram (EOG). We used four auxiliary electrodes (one located above and below the right eye, one located to the right of the left eye and right of the right eye). The Active two system band-pass filtered the EOG and EEG signals between 0.016 and 250 Hz prior to sampling. This also naturally limits the low frequencies we can reliably measure with this system. The signals were then digitally high-pass filtered (0.01 Hz cutoff) and notch filtered at 60 Hz to remove line noise. Robust PCA^85^ was used to remove artifacts and outliers from both EEG and EOG data, and subsequently both signals were digitally low-pass filtered (64 Hz cutoff) and down-sampled to 128 Hz. Bad EEG electrode channels were identified manually and replaced with interpolated channels. The interpolation was performed using the 3D cartesian coordinates from the electrode cap projected onto a plane using all surrounding “good” electrodes. Any faulty EOG channels were removed, e.g. if no signal was recorded or the power exceeded 3 standard deviations of the other channels. The EOG channels were then used to remove eye-movement artifacts by linearly regressing them from the EEG channels, i.e. least-squares noise cancellation. In each EEG channel, additional outlier samples were identified as values exceeding 4 times the distance between the 25th and the 75th quartile of the median-centered signal, and samples 40 ms before and after such outliers were replaced with interpolated samples using neighboring electrodes.

#### Recording and preprocessing of ECG

The ECG signal was recorded using a flat electrode placed below the left lumbar region (Left Leg / LL position) with a BioSemi Active Two system at a sampling frequency of 2048 Hz. The ECG signal was detrended using a high-pass filter (0.5 Hz cutoff) and subsequently notch filtered at 60 Hz to remove line noise. Peaks in the ECG corresponding to the R-waves were found using *findpeaks* (built-in matlab function). The instantaneous HR is computed for each beat as the inverse of time intervals between subsequent R-wave peaks and interpolated using cubic spline interpolation. To ensure the same sampling frequency for all subjects this instantaneous HR signal is resampled at a regular sampling rate of 128Hz.

#### Recording and preprocessing of respiration and respiratory volume

The respiration signal was recorded at a sampling frequency of 2048 Hz using the BioSemi Active Two system by a SleepSense 1387 Respiratory Effort Sensors, which captures the tension on a belt worn around the chest of the subject. The polarity of the signal was detected using peaks in the respiration signal (deep breaths) and inverted where necessary to ensure the correct phase of the signal. To compute the respiratory volume signal, we compute the envelope of the respiration signal by taking the absolute value of the hilbert transformed signal.

#### Intersubject correlation (ISC)

To determine whether a given factor drives responses we measured the temporal correlation of these signals between pairs of subjects. For most signals this is simply the Pearson’s correlation coefficient. For the scalp potentials (EEG) we use correlated-component analysis^86^.

We can compute the intersubject correlation for the gaze position, gaze variation, pupil size, heart rate, saccade rate, respiration, respiratory volume, electro-dermal activity signals by following 3 steps: 1. computing the Pearson’s correlation coefficient between a single subject’s signal (each of the modalities independently) and that of all other subjects while they listened to a story or carried out the control experiments. 2. a single ISC value for that subject was obtained by averaging the correlation values between that subject and all other subjects. 3. The two first steps are then repeated for all subjects, resulting in a single ISC value for each subject. For ISC of gaze position we compute the ISC in the horizontal and vertical gaze direction using the procedure as described above separately. To obtain one single ISC value we average the ISC for the horizontal and vertical directions.

The advantage of this technique is that we are not making any assumptions about which feature of the narrative drives these signals, but rather, only assume that the features we captured affect these signals similarly in different subjects. As such, we can capture potentially complex non-linear responses of the signals in questions to the various drivers, despite “only” measuring correlation. In contrast, the WSC and TRF approaches, which also measure correlation, are limited to first-order (linear) relationships between the two signals that are analyzed.

#### Frequency resolved intersubject correlation (synchronization spectrum)

We performed a frequency analysis to investigate at which time scale the recorded signals correlated between subjects. Each signal from each subject was band-pass filtered using 5th order Butterworth filters with logarithmic spaced center frequencies with a bandwidth of 0.2 of the center frequency. The ISC was computed for each subject in each frequency band for all videos. To obtain a single ISC value per frequency band we average ISC values for all videos and subjects.

#### Frequency resolved within subject correlation (WSC)

To capture the correlation between different signal modalities we measure the correlation between signals in different frequency bands (coherence). This is done within subjects, overcoming the limitation of ISC, which requires a common driver to synchronize signals between subjects. Thus, WSC can be measured in the absence of a common driving force such as during rest. But also, note that if there is a common driving force, WSC may measure a significant correlation that is due to this common factor, and not indicative of a direct causal link between the signals. To measure WSC signals are band-pass filtered (using the same filters as used for the frequency resolved ISC) and correlation is measured with real and imaginary components of the Hilbert transform capturing any potential time delays between signals.

We can do this procedure between one dimensional signals, but for neural signals we follow the procedure as in^33^. We utilize least squares regression between the EEG signals predicting each of the peripheral signals used in this study. These linear projections of this sort are conventionally referred to as components of the EEG. WSC is then Pearson’s correlation between the predicted signal (from the EEG signal) and the peripheral signal in question. In the cases where we differentiate between conditions, we compute component weights and WSC in each condition separately.

WSC in all instances was computed on test data using leave-one-subject out cross-validation (i.e. regression parameters are estimated including all subjects except for the test subject; WSC is then computed on the test subject; this train-test process is then repeated for all subjects.) Statistical significance is established in the same way as for ISC as described in previous sections.

#### Cluster statistics for contrast between conditions in frequency resolved correlation

To determine significant differences in spectra in different conditions e.g. between Attentive and distracted conditions (e.g. Fig. 2D and 3B), or control conditions and resting state (e.g. Fig. 5D and 5E) we use cluster shuffle statistics as follows. Since different frequency bands in the frequency-resolved ISC, WSC or SRC are not independent, we use one-dimensional cluster statistics including random shuffles to correct for multiple comparisons following an established procedure^87^. The procedure involves four steps: 1) take the difference between the spectra in two different conditions (e.g. Attentive and the distracted) for each subject. 2) compute a one-sample t-test on the difference for each frequency band. 3) clusters are identified as consecutive frequency bands with p-values below 0.01. The t-values within each cluster are then summed. 4) run 10,000 permutations in which we randomly change one half of the subjects sign of difference between the spectra computed in step 1. Steps 2-3 are then repeated while keeping the sum of t-values of the largest cluster. Finally, we compare the clusters’ t-values obtained in step 3 with the distribution of permuted cluster t-values obtained in step 4. Clusters with larger than 99 % (corresponding to p-value<0.01) of the permuted distribution were considered significant after multiple comparison cluster correction. Note that for the contrast in attention using the EEG data, all data was used for optimization (of components of the EEG) including attentive and distracted conditions, so that any difference can not be due to the optimization procedure.

#### Cluster statistics for significance of frequency resolved correlation

To determine if frequency-resolved ISC, SRC and WSC values are significantly different from zero we use a similar cluster statistic as above. For cases involving EEG (correlated component analysis and regression) we use test data to avoid upwards bias due to optimization, which is performed on separate training data. The shuffling and cluster correction procedure consists of the same 4 steps as above, except that we divide subjects in two equal size groups at random. The premise of this is that values around zero will not differ significantly if placed at random in two different groups^87^.

#### Temporal response function (TRF)

To capture the specific time delay of correlation between signals we use the classic least-squares linear systems-estimation of the impulse response, which convolves from one signal to predict the another. We allow for acausal delays to determine which signal lead and which signal follows, and estimate the filters in both directions (say x(t)=h_xy_(t) * y(t) and y(t)=h_yx_(t)* x(t), where * represents a convolution. Non-zero coefficient in positive lags of h_yx_(t) indicates that fluctuations in x(t) lead fluctuations in y(t), and vise versa, non-zero coefficient in positive lags of h_xy_(t) indicates that y(t) leads x(t). This is the basis for Granger “causality” ^88^. However, Granger causality is confounded if there are unobserved common factors. Since that is typically the case here, we refrain from using Granger causality as a tool, and simply report which signal is leading or following another.

#### Significance of temporal response function

To determine which time delays of correlation between signals are significant for the temporal response function we use shuffle statistics. 1) We compute the filter between the input and output signal (as described above). 2) We then repeat the procedure but now using a circular time-shifted version of the input signal. The delay is chosen randomly from a distribution with a maximum delay of the duration of the input signal. We repeat procedure 2 using 10.000 delays allowing us to discern p-values down to 0.001. To determine whether a specific time delay is significant we estimate the p-value for each time delay by comparing the filter estimated in step 1 to the surrogate distribution of filters found in step 2. I.e. how many of the surrogates had a value below the ones found in step 1. We correct the p-values found for each time delay using False Discovery Rate (FDR). We use a p-value of 0.01 to determine whether a specific time delay is significant.

#### Semantic speech features

To investigate which aspects of the speech signal our brain and body track, we construct semantic features of the speech signal. Previous work has shown that semantic dissimilarity produces a N400-like response in speech to speech signals using scalp recordings^38^. We follow the same approach here, by first transcribing and obtaining start/stop times for each word in the stories used in the study using a combination of Automatic Speech Recognition (ASR) and manual corrections. To compute the semantic dissimilarity we follow the same approach as^38^ we 1) quantize each word using the word2vec word embedding model (Matlab implemented version). 2) for each word we compute the correlation distance (one minus the Pearson correlation between the two word embedding vectors) between the present word and the average embedding vector of the preceding words in the sentence. This is done to capture the context of the sentence. For the first word in a sentence the dissimilarity is computed between the average word embedding of the preceding sentence, the first word in the story is ignored. The second semantic feature we use is the word rate as a measure of lexical difficulty. We compute the word rate as the inverse of the inter-word interval, which we then resample at regular sampling intervals.

## Footnotes

1 We do not mean to imply that the cortex is “responding” to changes in pupil size. We only use the terminology of TRF to stay consistent with the terminology that was used when linear systems modeling was introduced to EEG analysis^54, 55^.

