## Supplement for "Narratives engage brain and body: bidirectional interactions during natural story listening"

### Supplemental information for “Narratives engage brain and body: bidirectional interactions during natural story listening”

Jens Madsen, Lucas C. Parra
Department of Biomedical Engineering, City College of New York
85 St. Nicholas Terrace, New York, NY 10031, USA

**Section S1: Effects of blinks and saccades on pupil size during story listening**
When we blink our eyes, the pupil contracts briefly to protect the iris from potential harm. During eye tracking, when we blink there is a brief period of time where the pupil is partially occluded both before and after the blink, leaving the estimation of the pupil erroneous. These two effects are rather well known in pupillometry research and standard steps of preprocessing are used to account for this (+-100ms before and after a blink is removed and the entire blink is interpolated). However these are not the only effects that can affect the pupil size. Recent work has shown longer effects of pupil size in the order of seconds[1]. This made us investigate whether this effect had any cognitive effect and to what extent this happens during natural story listening. We first simply take the average of the change in pupil size (as compared to baseline) in a period of 4 seconds before and after a blink (Fig. S1A). Here we see a change in pupil size both before and after blinks. To verify which period of time we find a significant change in pupil size we compute the Temporal Response Function (TRF) between blinks and pupil size (Figure S1B). We find a significant change in pupil size due to blinks from 2 seconds before until about 5 seconds after the blink. We cannot rule out a common drive that elicits both blinks and changes in pupil size, so these TRFs do not necessarily reflect a causal effect of the blink on pupil size. However, the effect is present both in the Attentive and Distracted conditions, suggesting this is a low level effect of blinks on pupil size. As a result, this blink-pupil interaction could leave measurements of pupil size to be a secondary effect to blinks. We therefore remove the effect using traditional linear regression also referred to as “noise cancellation”. Essentially we estimate a TRF predicting the pupil size using blinks as input signal on the entire recording of the experiment. We verify that we have removed any linear effects of the blinks on the pupil size by computing the TRF on this new regressed signal (Fig. S1C).

Blinks are not the only potential confound that could cause a change in pupil size, saccades which are rapid movements of our eyes also cause a change in pupil size.[2] During the experiment people did look at a gray screen and all instructions on the screen were isoluminant. Despite this, we see a change in pupil size both before and after a saccade (Fig. S1D). We quantify the effect the saccades have on the pupil size using again the formalism of TRFs. We find a significant effect of saccades on pupil size from 4 seconds before until 4 seconds after the saccade. As with blinks we cannot rule out any common drive of pupil size and saccade initiation. Since we are specifically looking for cognitive effects of pupil size and the effect we see here is not modulated by attention. We therefore attribute this to a low-level saccade-pupil interaction and remove this effect using TRF regression, which we confirm by repeating the process after regressing out the saccade effect. The analysis of pupil size in Fig. 3 of the main uses this pupil size signal after effects of blinks and saccade have been jointly regressed out from the pupil size signals.

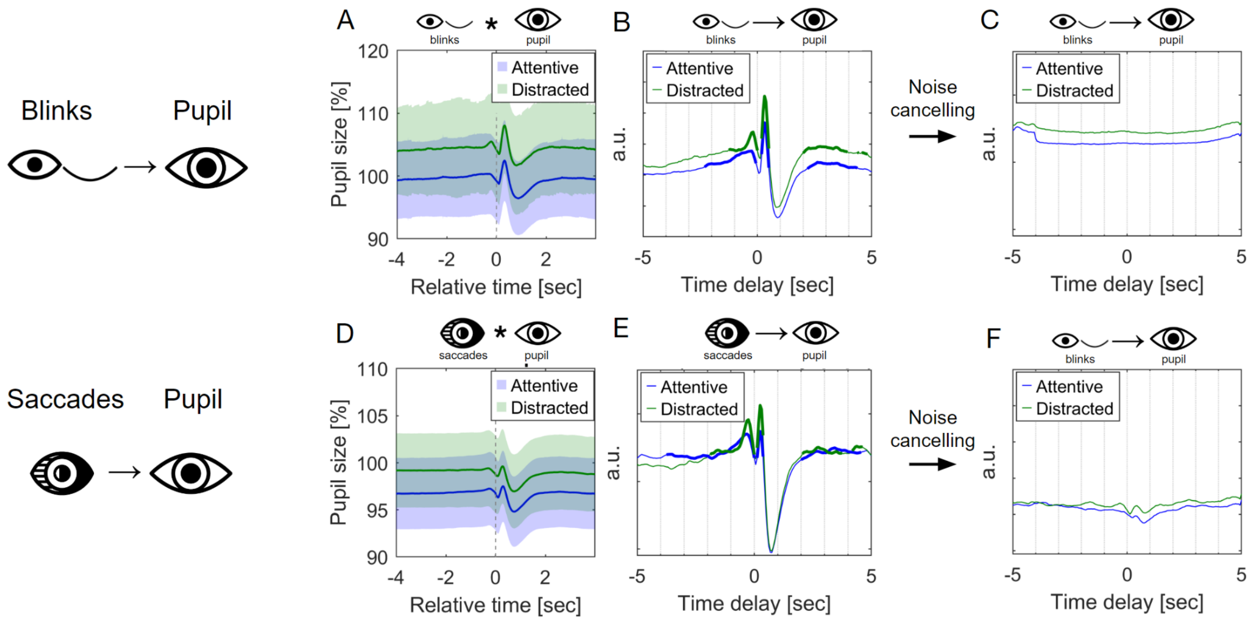


**Figure S1. Effects of saccade and blinks on pupil size during story listening. A)** Blink evoked pupillary response is the change in pupil size, relative to baseline, as a function of time relative to the offset of a blink (reverse correlation). The shaded area represents the SEM across participants (N=29 subjects). **B)** Temporal Response functions using pupil size as output signal and blinks as input (the blink signal is set to 1 whenever a participant is blinking and otherwise 0). Bold line indicates a significant difference in TRF between conditions computed using shuffle statistics with FDR correction (p<0.01, N=29 subjects). **C)** Temporal Response functions using pupil size (after saccades and blinks have been regressed out using the combined TRF of blinks and saccades) as output signal and saccade offsets as input signal. **D)** Saccade evoked pupillary response is the change in pupil size, relative to baseline, as a function of time relative to the offset of a saccade (reverse correlation). The shaded area represents the SEM across participants (N=29 subjects). **E)** Temporal Response functions using pupil size as output signal and saccades as input (the saccade signal set to 1 whenever a participant is saccading and otherwise 0). Bold line indicates a significant difference in TRF between conditions computed using shuffle statistics with FDR correction (p<0.01, N=29 subjects). **F)** Temporal Response functions using pupil size (after saccades and blinks have been regressed out using the combined TRF of blinks and saccades) as output signal and blink offsets as input signal.

**Section S2: Timing of saccade initiation during story listening is modulated by attention**

We found that there is a higher likelihood of saccade initiation at times of word onsets and this effect is modulated by attention (Fig. 2B). Given this finding, it is possible that people saccade more during attentive listening as compared to distracted listening.

*Number of saccades during story listening*

In Experiment 1 where people listened to the stories while looking at a blank gray screen, we did not find a different number of saccades during attentive listening (202.07 +- 180.26 saccades) as compared to distracted listening (199.79 +- 122.20 saccades). We specifically test this using a 3-way anova using subjects as a random effect, stimuli and attention as fixed effects. We find a significant effect of subjects $F(28,541)=9.68$, $p=1.31e-32$, a significant effect of stimuli $F(9,541)=16.79$, $p=1.72e-24$ (this is to be expected as the stories are of different length), however no main effect of attention $F(1,541)=0.05$, $p=8.18e-01$. This is however not replicated for our second experiment where in the attentive listening condition (168.34 +- 91.86 saccades) we find that people saccade less as compared to the distracted condition (185.93 +- 102.01 saccades). This is confirmed using a 3-way anova using subjects as random effect. We find a main effect of subject $F(37,694)=22.65$, $p=3.67e-95$, stimuli $F(9,694)=59.95$, $p=8.11e-81$ and a significant effect of attention $F(1,694)=28.37$, $p=1.36e-07$. This is perhaps expected as subjects are instructed to focus their gaze on an isoluminance fixation cross, which is itself a task requiring some level of attention.

*Size of saccades during story listening*

We found that people had larger gaze variation during attentive listening as compared to when they were distracted. Despite those fluctuations being at a much slower timescale than saccades, we wanted to test if people also had a difference in the size of their saccades during listening. In Experiment 1 during attentive listening we find people saccades in average 3.35 ± 1.45 Visual Degree Angle (VDA) as compared to 3.42 +- 1.80 VDA when they were distracted. We test the significance of this difference using a 3-way anova with the subject as a random effect. We find a significant subject effect ($F(28,533)=14.75$, $p=6.80e-50$), no significant stimuli effect ($F(9,533)=1.05$, $p=0.40$) and no attention effect ($F(1,533)=0.57$, $p=4.50e-01$). We repeat the analysis for Experiment 2 for attentive listening (2.72 ± 1.88 VDA) and distracted listening (3.06 ± 3.52 VDA) where we find a significant main effect of Subject ($F(37,702)=12.90$, $p=4.46e-57$), no significant stimuli effect ($F(9,702)=0.98$, $p=0.45$) and a borderline attention effect ($F(1,702)=4.21$, $p=0.04$).

*Saccade rate fluctuations*

We previously reported that saccade rate fluctuations in the course of an audiovisual narrative, and that these fluctuations are synchronized across subjects, provided they are paying attention.[3,4] We see similar fluctuations in saccade rate here during audio-only narratives (Fig. S2E). We also see that saccade rate is synchronized between subjects at a slow time scale (Fig S2F). However, contrary to the audiovisual narratives, this effect is not modulated by attention.

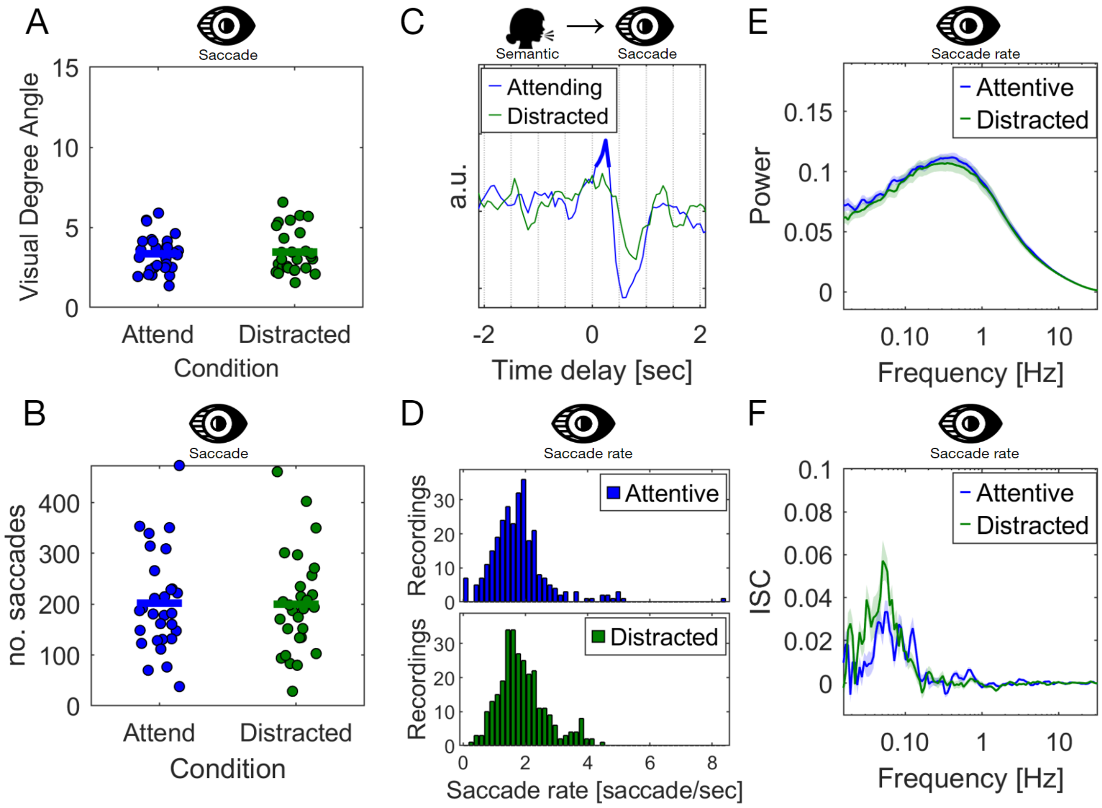


**Figure S2. Saccadic behavior during story listening. A)** Average size of saccades as measured by the Visual Degree Angle (VDA) when participants listened to 10 auditory narratives in an either Attentive or Distracted condition (each dot is a subject). Significance between the two conditions was established using a t-test ($p<0.05$). **B)** Number of saccades during story listening was computed as the average number of saccades across stories in the Attentive and Distracted condition (each dot is a subject, N=29). **C)** Temporal response function predicting saccade onsets from word onset timing. THe bold area indicates a significant temporal response estimated using circular shuffle statistics (see Methods). **D)** Histogram of saccade rate in the two listening conditions (Attentive and Distracted). The histogram is over the average saccade rate for each subject when they listened to each story. **E)** Power spectrum of the instantaneous saccade rate, which is computed as the inverse of the inter-saccadic interval, is computed by first taking the standard deviation for each subject and taking the mean over stories and vertical/horizontal direction and lastly normalized by the total power per subject across conditions. Line indicates mean over subjects and shading is the SEM across N=29 subjects. Significance of the power between the attentive and distracted condition is established in each band using Wilcoxon rank sum test, corrected for multiple comparisons using one-dimensional cluster statistics (light gray area indicates $p<0.01$, cluster corrected). **F)** Frequency-resolved ISC of saccade rate during story listening computed in similar fashion as Panel D.

**Section S3: Timing of blinking during story listening is modulated by attention**

We found that blink onset was predictable from word onsets and this effect was modulated by attention (Fig 2D). Based on this finding we wanted to know if people would blink more or less during the attentive condition compared to the distracted condition.

*Number of blinks during story listening*

We expect there to be a stimuli effect as the duration of each story is different, we would also expect there to be a subject effect. For Experiment 1 in the attentive listening condition we found people blinked (90.99 ± 159.37 blinks) as compared to 91.74 ± 80.34 blinks in the distracted condition. We test the difference using a 3-way anova and find a significant main random effect of subjects ($F(28,541)=5.71$, $p=1.95e-17$), an expected stimuli ($F(9,541)=5.02$, $p=1.61e-06$) and no effect of attention ($F(1,541)=0.01$, $p=0.93$). For Experiment 2, with a fixation cross, we did find a difference in the number of blinks between the attentive condition (97.93 ± 58.23 blinks) as compared to the Distracted condition (105.82 ± 72.54 blinks). We test this again using a 3-way anova where we confirm the attention effect of ($F(1,694)=16.74$, $p=4.79e-05$).

*Duration of blinks during story listening*

We also found that the timing of when blinks occur during story listening was modulated by attention (Fig. 2C). We distract people from the story by asking them to carry out a mental arithmetic task whereby they silently count backwards from a high prime number (between 800-1000) in decrements of 7. In this dual task we speculated that the duration of blinks could be longer, essentially allowing for internal cognitive processing of the counting. This is indeed what we find, during the attentive listening peoples blink duration is on average 0.23 ± 0.50 seconds as compared to 0.40 ± 1.25 seconds during the distracted condition. We confirm the difference in blinking duration using a 3-way anova with subject as a random effect ($F(28,533)=1.37$, $p=9.75e-02$), stimuli and attention as fixed effects. We find a significant attention effect ($F(1,533)=5.02$, $p=0.03$). We repeat these findings for Experiment 2. In this attentive listening condition (blink duration: 0.24 ± 0.47 sec) blinks are shorter as compared to when they are distracted (blink duration: 0.32 ± 0.62 sec). We confirm this observation using the same 3-way anova. We find a significant effect of attention $F(1,683)=5.98$, $p=0.01$.

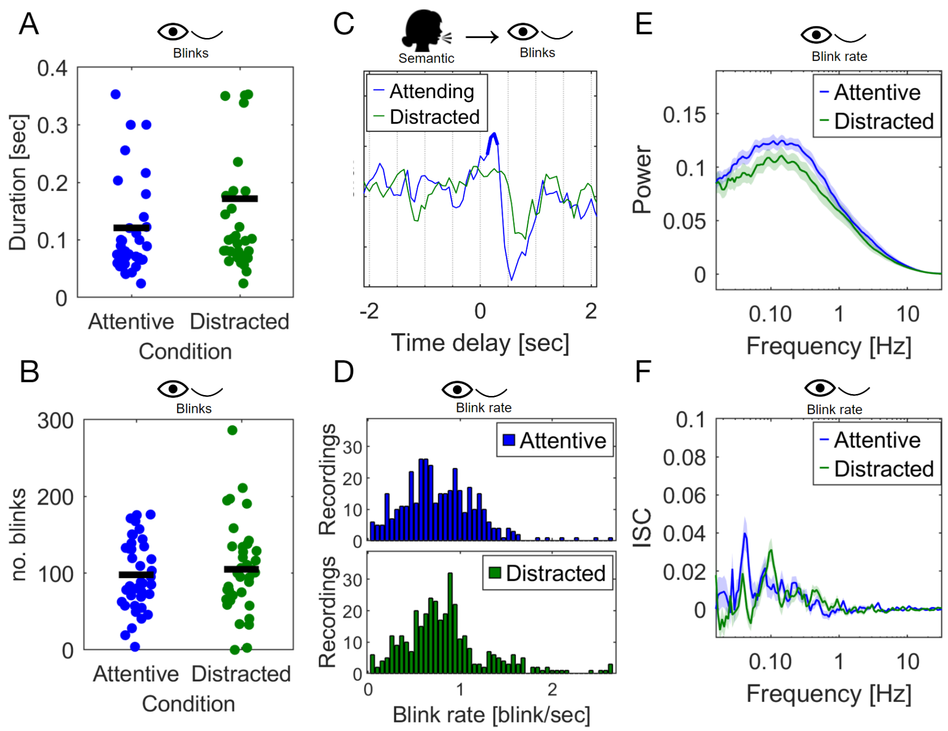


**Figure S3. Blinks during story listening. A)** Average duration of blinks when participants listened to 10 stories in an either Attentive or Distracted condition (each dot is a subject). Significance between the two conditions was estimated using a t-test ($p<0.05$). **B)** Average number of saccades across stories when participants listened to the stories in an Attentive or Distracted condition (each dot is a subject, N=29). **C)** Temporal Response Function using word onsets as input signal and blinks as output signal (the blink signal was set to 1 whenever a participant was blinking and otherwise 0). **D)** Histogram of blink rates accumulated across participants and stories as they listened to 10 stories in an Attentive and Distracted condition. The blink rate is computed as the instantaneous blink rate averaged across each story for each participant. **E)** Power spectrum of the instantaneous blink rate, computed as the inverse of the inter-blink intervals, is computed by first taking the standard deviation for each subject and taking the mean over stories and vertical/horizontal direction and lastly normalized by the total power per subject across conditions. Line indicates mean over subjects and shading is the SEM across N=29 subjects. Significance of the power between the attentive and distracted condition is established in each band using Wilcoxon rank sum test, corrected for multiple comparisons using one-dimensional cluster statistics (light gray area indicates $p<0.01$, cluster corrected). **F)** Frequency-resolved ISC of blink rate during story listening computed in similar fashion as Panel B.

**Section S4: Brain-body interaction of eye behavior and scalp potentials during story listening**

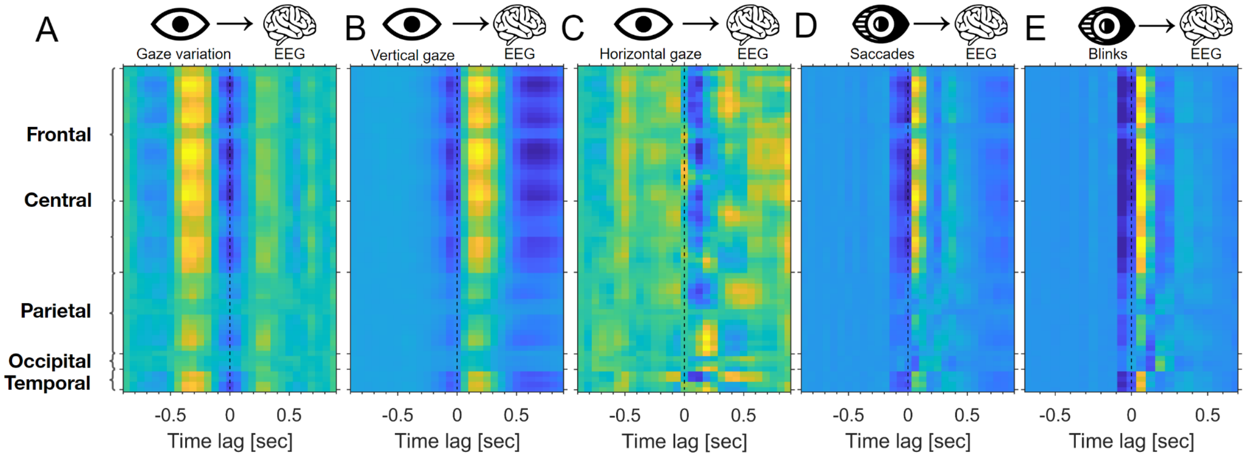


**Figure S4 Brain-body interaction between eye behavior and scalp potentials during story listening. A)** Temporal Response Function between predicting scalp potentials from Gaze variation. Gaze variation is computed as the hilbert transformed

**Section S5: Eye movements during story listening**

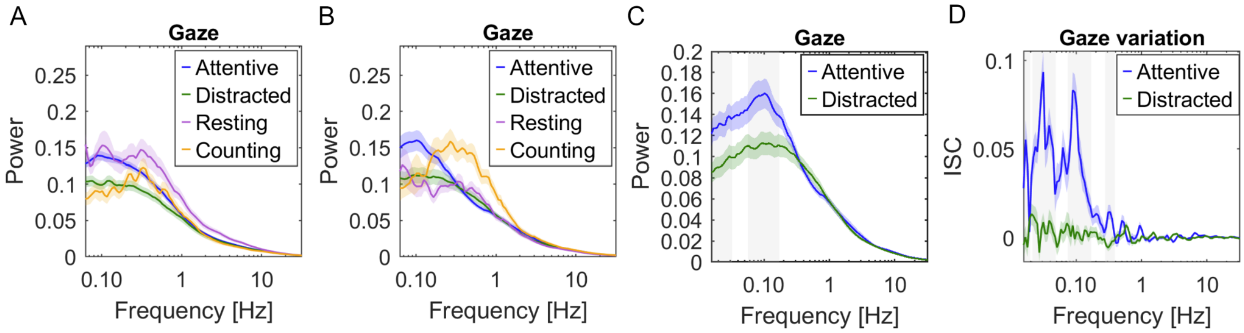


**Figure S5: Eye movements during story listening. A)** Power spectrum of gaze position during attentive listening, distracted listening, resting state and counting condition for Experiment 1 - no fixation cross (N=29). The power spectrum was computed as the pooled standard deviation for gaze position in the vertical and horizontal directions in each frequency bin. The shaded area is the standard error of the mean. **B)** same as panel B but for Experiment 2 (N=38) where a fixation cross was shown on the screen for all conditions. **C)** Power spectrum of gaze position for Experiment 2 (N=38) while participants listened to the 10 auditory narratives in the attending and distracted conditions. **D)** The synchronization spectrum of gaze variation for Experiment 2 (N=38). Gaze variation is computed as the absolute value of the Hilbert transformed gaze position signal.

**Figure S6: The relationship between speech features and pupil size during story listening**

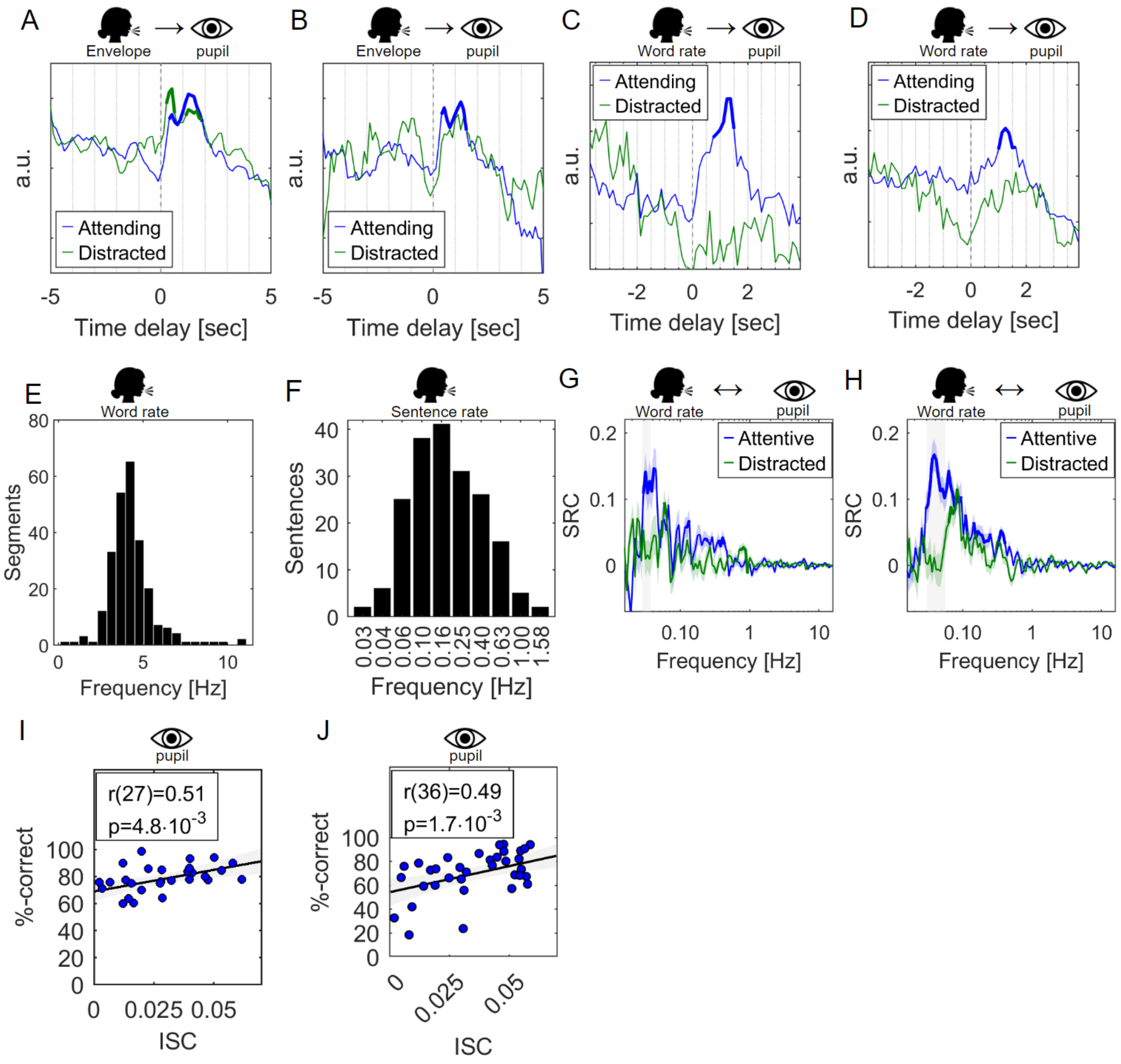


**Figure S6: Temporal response function between speech features and pupil size during story listening. A)** Temporal response function using the speech envelope as input signal, computed as the absolute value of the hilbert transformed speech signal. The output signal is the pupil size as subjects (Experiment 1, N=29) listened to the 10 stories (Duration 27 minutes) )included in the experiment. **B)** Temporal response function computed the same way as panel A but for Experiment 2 (N=38). **C)** Temporal response function using word rate (words spoken in the narrative per minute) as input signal predicting the fluctuations of pupil size (Experiment 1). **D)** same as panel C but for Experiment 2. **E)** Power spectrum of the word rate for the 10 stories used in the study. The grayed area is the standard error around the mean (N=10 stories). **F)** Stimulus Response Correlation (SRC) between the word rate of the stories people listened to and the pupil size of participants as they listened to the narratives in an attending (blue) and distracted condition (green). Bold line indicates significant correlation computed using t-test corrected for multiple comparisons using cluster correction (p<0.01). Grayed area indicates significant difference between the attending and distracted conditions using the same technique as the previous approach (p<0.01). **G)** same as panel F but for Experiment 2. **I)** Intersubject correlation of pupil size (bandpassed between 0.01Hz and 1Hz) and the score subjects got from 4 alternative forced choice questionnaires they received after the stories were played (Experiment 1). Each is the average of each participant's score on all questions and the average ISC across the 10 stories they listened to in the attentive condition. The correlation between the score and ISC was pearson correlation. **J)** same as panel I but for Experiment 2.

**Figure S7: Electro-dermal activity and respiration behavior during story listening**

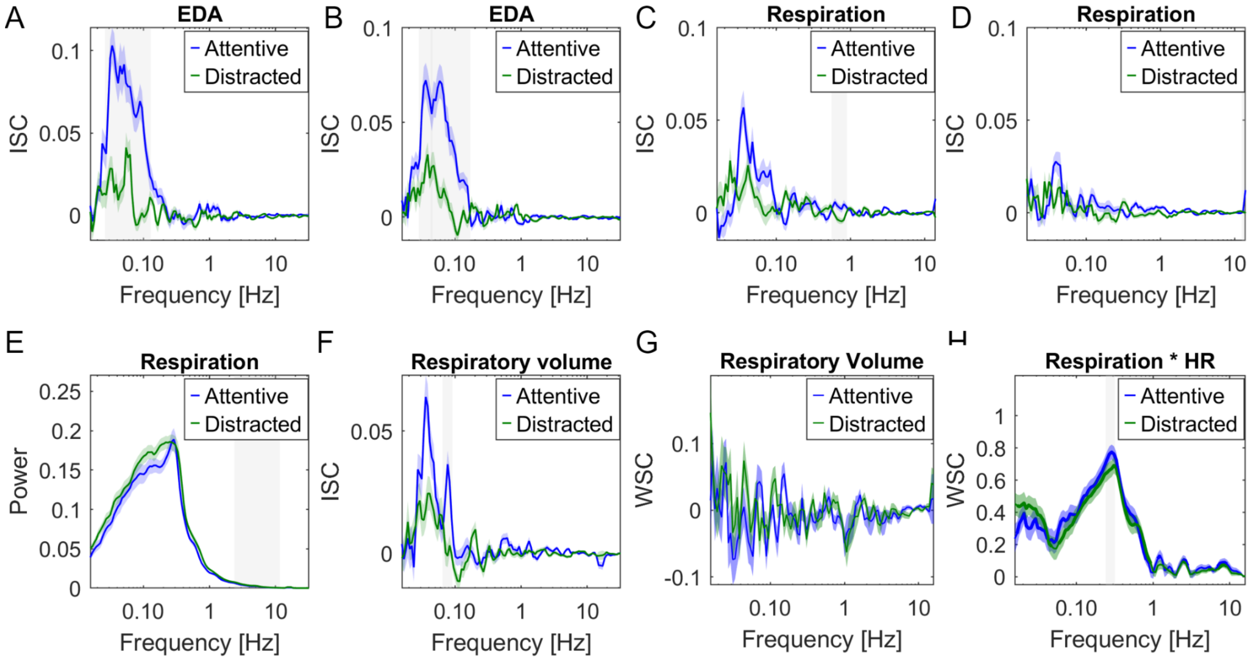


**Figure S7. A)** Synchronization spectrum of Electro-Dermal Activity (EDA) while participants listened to auditory narratives in an Attending (blue) or Distracted condition (green) for Experiment 1. Light gray shading indicates a significant difference between the attentive and distracted conditions using a t-test corrected for multiple comparisons using one-dimensional cluster statistics (p<0.01, cluster corrected, N=29) **B)** same as panel A but for Experiment 2. **C)** Synchronization spectrum of Respiration for Experiment 1 during the auditory narratives. **D)** same as panel C but for Experiment 2 (N=38). **E)** Power spectrum for respiration during auditory narratives for Experiment 1. **F)** Synchronization spectrum of respiratory volume for Experiment 1 (N=29). **G)** Within subject correlation computed between scalp recordings (EEG) and the Respiratory volume signal when people listened to the auditory narratives. **H)** Within subject correlation between Respiration and heart rate for Experiment 2 while participants listened to auditory narratives.

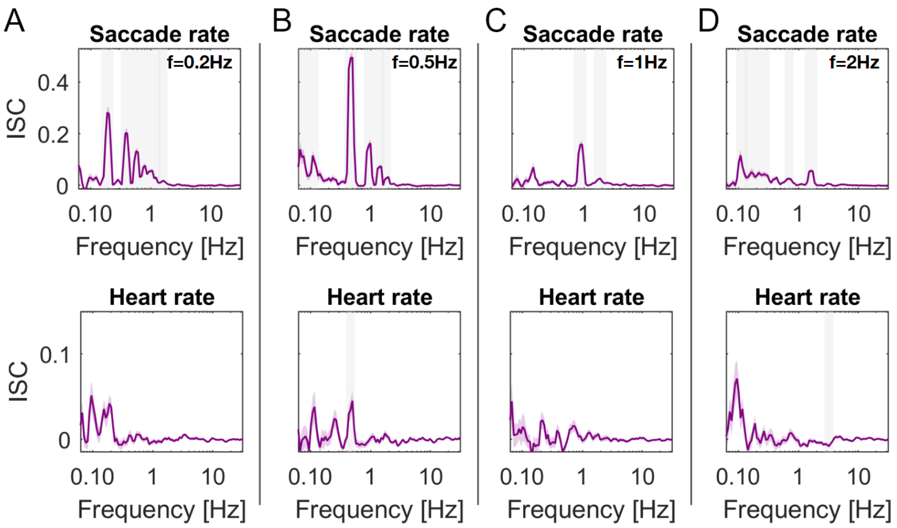


**Figure S8. Controlled saccades modulate heart rate. A)** Synchronization spectrum of saccade rate (upper panel) and heart rate (lower panel) during controlled saccade condition during Experiment 2 (N=38). People were asked to follow a dot jumping around on the screen at a rate of 0.2Hz, i.e. dots were displayed 5 seconds at a time. The bold line is the average across N=38 subjects and shading is the SEM. Light gray shading indicates a significant difference from 0 corrected for multiple comparisons using one-dimensional cluster statistics (p<0.01, cluster corrected) **B)** same as panel A but for a dot jumping at a rate of 0.5Hz. **C)** same as panel A but for a dot jumping at a rate of 0.5Hz. **D)** same as panel A but for a dot jumping at a rate of 0.5Hz.

**Heart-beat Evoked Responses during rest and attentive listening**

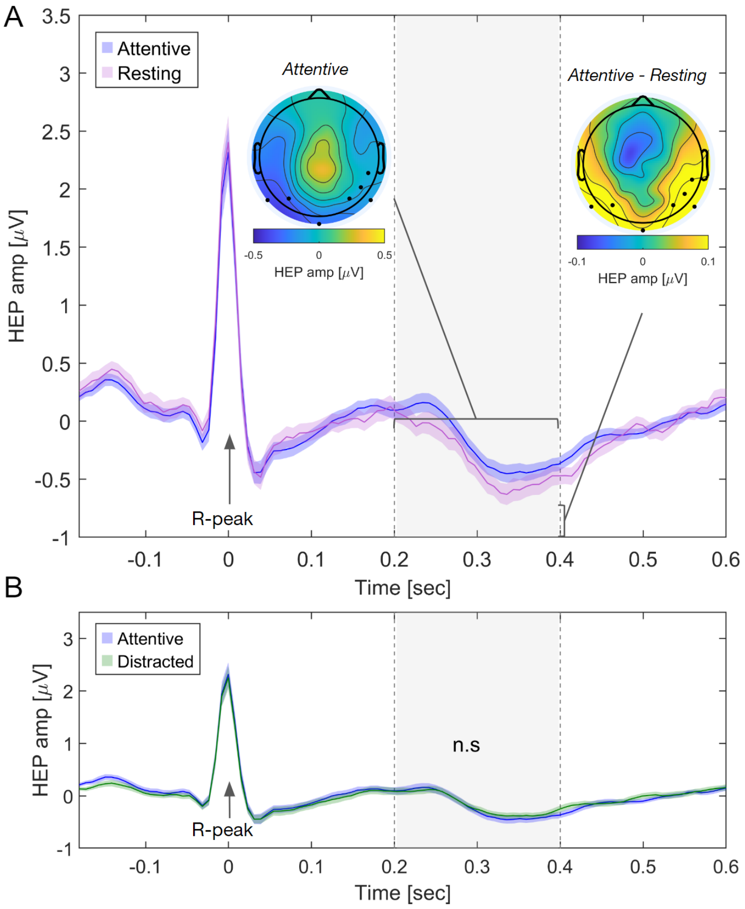


**Figure S9. Heart-beat Evoked Responses during rest and attentive listening. A)** Heartbeat Evoked Potentials (HEP) during the attentive listening and resting state conditions for Experiment 1. Scalp potentials were time-aligned to R-peaks of the ECG signals, averaged over the 10 stories. We averaged HEP in the range of 0.2-0.4 sec (gray shaded) based on prior literature showing cognitive effects in this time window. This average HEP is significantly enhanced in lateral posterior electrodes during attentive listening to the narrative as compared to rest (t-test across N=29 subjects, FDR corrected at 0.01, black dots in scalp distributions). The time course is the HEP averaged across this set of significant electrodes (selected post hoc and should thus be viewed with caution). Color-shaded areas indicate standard error of the mean across subjects. **B)** HEP during attentive and distracted listening conditions. No significant electrodes were found using the same method as panel A. The time course is the HEP averaged across the set of significant electrodes found as described in panel A.
